# Structural modification of oxazolidinone antibiotics alters nascent peptide stalling preference and peptide trajectory through the ribosome

**DOI:** 10.64898/2026.02.16.706219

**Authors:** Jordan I. Kleinman, Tushar Raskar, Dorota Klepacki, Teresa Szal, Nora Vázquez-Laslop, Alexander S. Mankin, James S. Fraser, Danica Galonić Fujimori

## Abstract

The oxazolidinone antibiotic linezolid binds to the peptidyl transferase center of the ribosome, where it inhibits a subset of peptide bond formation events. This context-specificity of translation inhibition is dictated by the nature of the amino acid at the penultimate position of the nascent peptide. It remains unknown whether this is a general feature of oxazolidinones and whether it can be modulated by their structural alterations. Here, we show that the oxazolidinone tedizolid also inhibits translation in a context-specific manner, but with dramatically altered selectivity, favoring Ile, His, and Gln as the penultimate residues. Delpazolid, which shares the C5 hydroxymethyl moiety with tedizolid, shows a similar preference. Structural analysis of the ribosome with tedizolid and a stalled nascent peptide showed a compacted, helical conformation of the nascent chain induced by the drug. Our findings reveal that stalling preferences of oxazolidinones can be modulated by structural modifications within this antibiotic class.

## INTRODUCTION

The peptidyl transferase center of the ribosome (PTC), situated within the large ribosomal subunit, catalyzes peptide bond formation. Within the PTC, peptide bond formation occurs between the C-terminal amino acid of the nascent chain, tethered to the P-site tRNA (P-tRNA), and the incoming amino acid delivered in the A site by aminoacyl tRNA (aa-tRNA). Stabilization of the PTC–proximal region of the nascent peptide in a specific, β-strand–like conformation orients the donor substrate for efficient catalysis of peptide bond formation^1^.

Ribosome-targeting antibiotics interfere with various stages of translation ^2,3^. PTC– and ribosome exit tunnel-binding antibiotics have long been considered global inhibitors of translation, which interfere with peptide bond formation by preventing proper accommodation of PTC substrates or by plugging the exit tunnel, respectively^4–6^. Recent investigations have shown, however, that several among these antibiotics exhibit nascent peptide-dependent inhibition of translation. Rather than indiscriminately stalling the ribosome, the effect of these drugs varies depending on the sequence of the nascent polypeptide chain^4–6^. Instead, these compounds inhibit translation more efficiently when the ribosome encounters specific sequences of amino acids in the nascent chain. This phenomenon of context-specific inhibition of translation has been documented for numerous exit tunnel-binding antibiotics, including a variety of macrolide antibiotics^7,8^ and polyketide tetracenomycin X^9^, for the PTC targeting inhibitors, such as chloramphenicol and linezolid^10–14^, as well as for other ribosome-binding antibiotics^15–17^.

While detailed analysis of specific antibiotic-dependent stalling sequences has been made possible by structural evaluation of drug-arrested translation complexes^9,11,12,18–22^, we lack a broader understanding of the mechanisms of context-specificity. More recent structural data points to a combination of effects by which stalling may occur in the presence of antibiotics, including allosteric modulation of critical PTC nucleotides and forcing PTC substrates into conformations which preclude successful peptide bond formation, depending on antibiotic identity and sequence of the synthesized polypeptide. A deeper understanding of the interplay between these mechanisms and how they are affected by structural differences within the antibiotic requires further exploration. Such studies would greatly benefit from antibiotics with related chemical structures but different context specificity of action.

The oxazolidinone class of antibiotics^23^ provides an ideal starting point for exploring the structural dependence of context-specific stalling (Fig. 1A). Linezolid^24,25^ (LZD), the first approved oxazolidinone antibiotic, and tedizolid^26^ (TZD), a more recently developed derivative, are clinically used for the treatment of Gram-positive bacterial infections. A related oxazolidinone derivative, radezolid^27,28^ (RZD), is in clinical development for the treatment of bacterial acne and community-acquired pneumonia. While distinct in their C and D rings, LZD and RZD share the C5 acetamidomethyl functionality. Both LZD and RZD inhibit translation in a context-dependent manner, with Ala in the penultimate position of the nascent peptide [Ala(−1)] as the critical sequence determinant. Cryo-EM analysis of LZD– and RZD-stalled ribosome complexes revealed the critical importance of the C5 acetamidomethyl group of these antibiotics in stabilizing antibiotic binding to the ribosome and the formation of a peptide binding pocket complementary in shape to the Ala(−1). Interestingly, although early structure-activity relationship analysis indicated the necessity of the C5 acetamidomethyl group for bacterial growth inhibition^24^, TZD lacks this functional group and, instead, carries a C5 hydroxymethyl substituent. The critical role of the C5 moiety of LZD and RZD in context-dependent stalling prompted us to investigate how replacement of this group with a smaller hydroxymethyl substituent affects translation inhibition.

**Figure 1:**
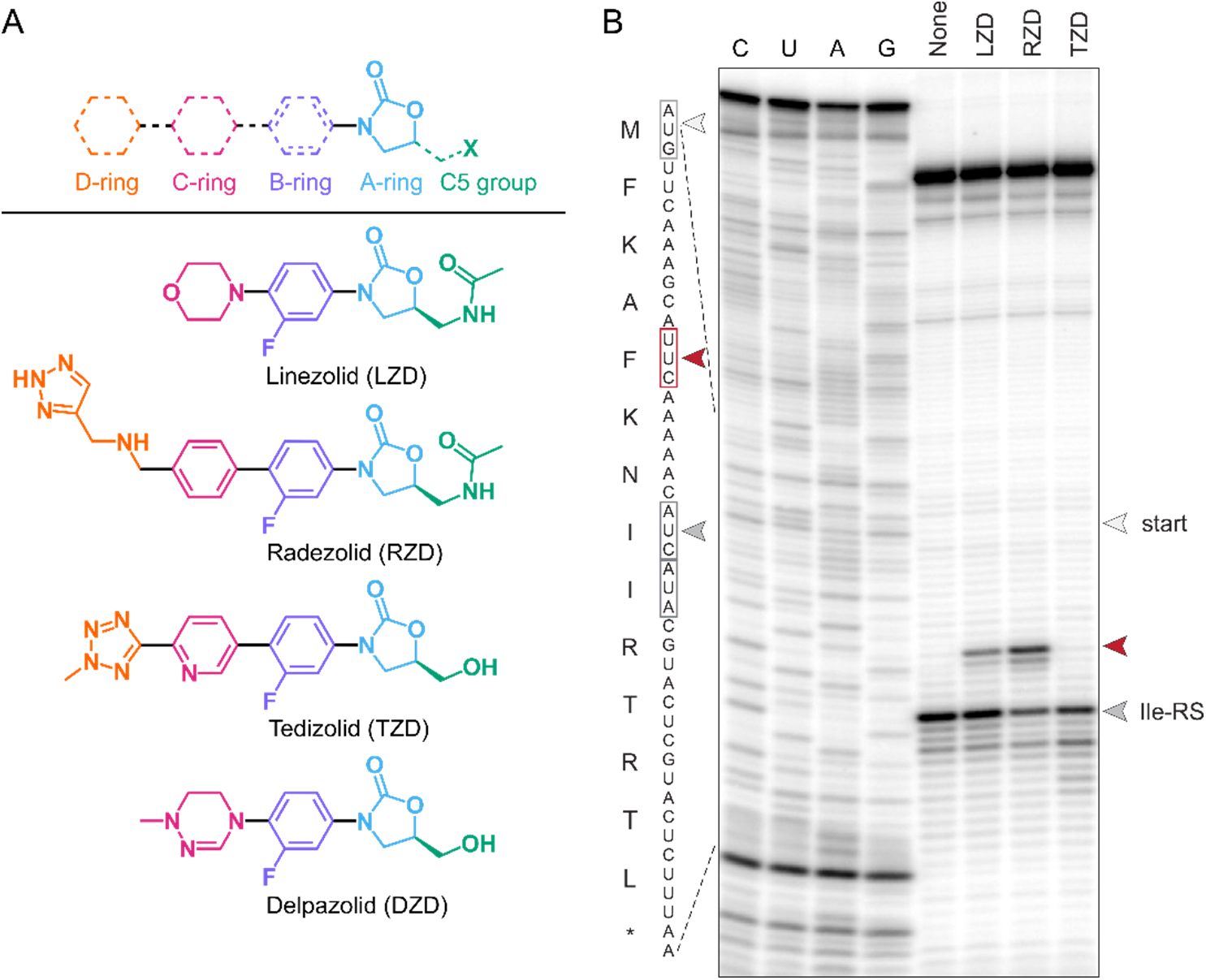
Tedizolid exhibits altered context-specificity compared to linezolid and radezolid. **A)** Structures of oxazolidinone antibiotics. Oxazolidinone antibiotics are composed of a modular multi-ring structure, with variation at the C/D rings and the oxazolidinone C5 substituent. **B)** Ribosome toeprinting performed on a model stalling mRNA in the presence of DMSO, LZD (50uM), RZD (10uM), or TZD (25uM). Stalling with Ala(−1) is indicated in red. Translation beyond the expected stall site is inhibited by isoleucyl-tRNA synthetase inhibitor mupirocin (50uM) present in all reactions (dark gray).

Here, we use *in vivo* ribosome profiling and *in vitro* toeprinting to show that TZD, as well as an investigational oxazolidinone delpazolid^29^ (DZD, Fig. 1A), which also contains the C5 hydroxymethyl group, exhibits context-specificity different from that of LZD and RZD. TZD and DZD preferentially stall the ribosome when isoleucine or histidine appear in the penultimate (−1) position of the nascent chain. Cryo-EM analysis of a 2.16 Å TZD-stalled ribosome complex revealed a broader peptide binding pocket created by the C5 hydroxymethyl compared to C5 acetamidomethyl, favoring larger amino acids at the –1 position. Strikingly, this wider pocket allows the nascent peptide to adopt a compact, helical conformation in the presence of TZD. This novel peptide path stabilizes multiple rRNA nucleotides in orientations that disfavor peptide bond formation. Our findings support an allosteric mechanism of drug-induced stalling by tedizolid, in which stabilization of an unusually compacted peptide conformation inhibits movement of critical rRNA nucleotides, thereby preventing continued translation.

## RESULTS

### Tedizolid exhibits altered context-specificity compared to linezolid and radezolid

To explore whether the stalling preference of TZD matches that of LZD and RZD, we performed ribosome toeprinting to examine the effect of this antibiotic on *in vitro* synthesis of a peptide containing the MFKAF stalling motif (Fig. 1B). It was previously shown that both LZD and RZD stall translation of this model peptide when Ala reaches the –1 position of the nascent chain^10,11,13^. *In vitro* transcription-translation was performed using *E. coli* ribosomes, in the presence or absence of drug. Ile-tRNA synthetase inhibitor mupirocin was included in all reactions to “catch” any ribosomes that progressed beyond the anticipated stall site.

Treatment with LZD or RZD resulted in a buildup of truncated product corresponding to Ala(−1) stalling, consistent with previous profiling^10^ and structural^11^ data. Surprisingly, no stalling was observed at this site in the presence of TZD. Instead, translation continued until the ribosome reached the “catch” codon. Thus, TZD appears to act in a context-specific manner distinct from that of LZD and RZD.

### Ribosome profiling reveals altered specificity of tedizolid-induced ribosome stalling

We used ribosome profiling as an unbiased strategy to identify stalling preferences of TZD. Oxazolidinone-susceptible *E. coli* strain BW25113 *ΔacrB* was treated with an excess of TZD (10x minimum inhibitory concentration) for 2.5 min, then rapidly flash-frozen and mechanically lysed. Ribo-seq and RNA-seq libraries were prepared as previously described^30^, and genes with >20% non-zero reads and an average read count of 0.5 across the coding sequence (CDS) were considered for downstream analysis.

Metagene analysis of the Ribo-seq reads mapped to the *E. coli* genome showed a clear 3-nt periodicity, attesting to the high quality of the Ribo-seq data^30^, whereas as expected, RNA-seq reads remained codon-independent (Extended Data Fig. 1A). Both Ribo-seq and RNA-seq datasets highly correlated between replicates (Extended Data Fig. 1B). Compared to DMSO control, TZD treatment led to a marked increase in ribosome density within the first ∼30% of the open reading frames (ORFs) transcriptome-wide, and a concomitant decrease in read density in the latter half of the ORFs (Fig. 2A), consistent with the drug acting as elongation inhibitor^31,32^.

**Figure 2:**
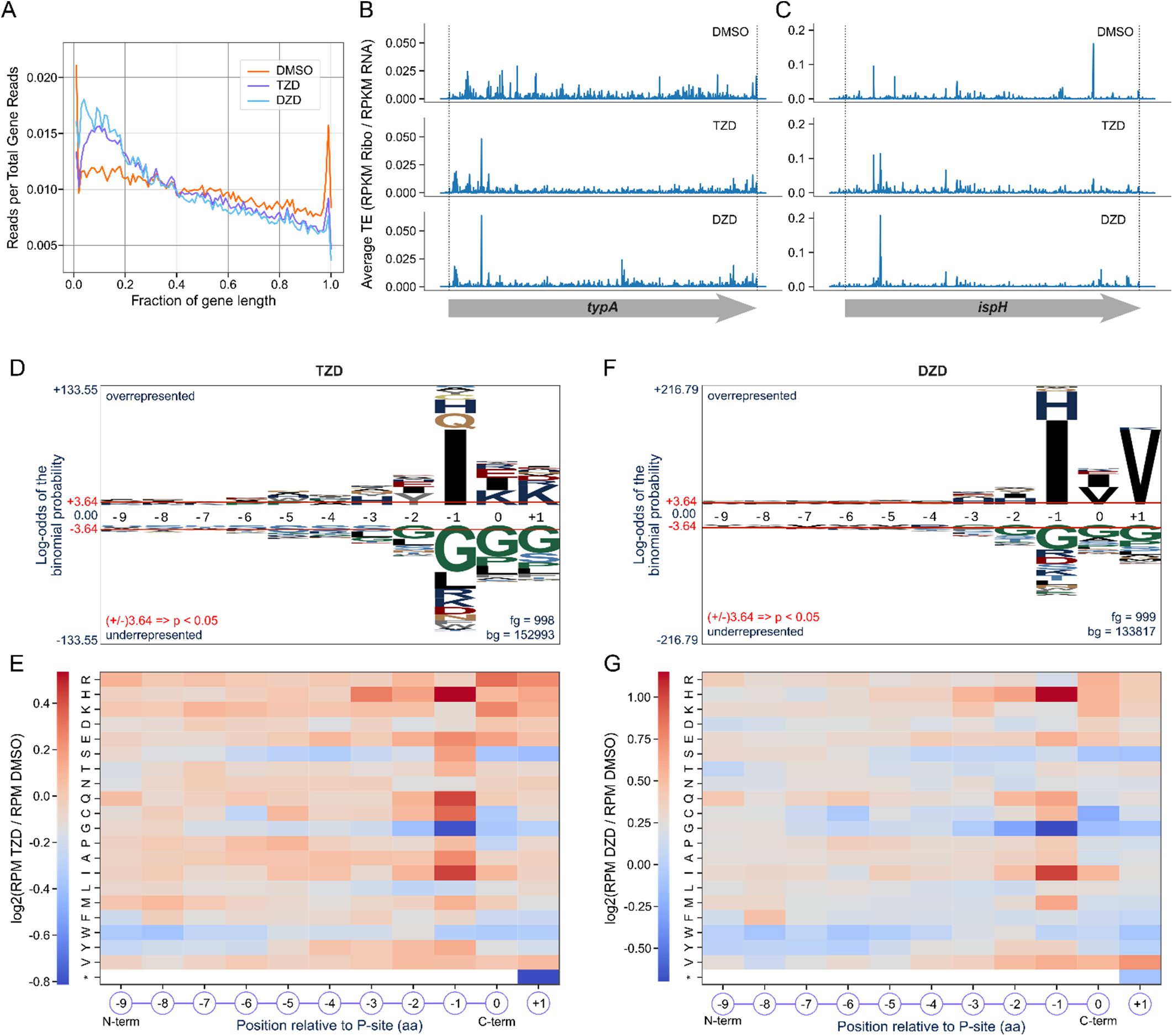
Ribosome profiling reveals unique stalling preferences of tedizolid and delpazolid. **A)** Metagene analysis of ribosome profiling data following 2.5 min treatment with 10x MIC TZD (20 μg/mL) or DZD (80 μg/mL). Positions within coding regions were normalized to gene length, and reads at each position were averaged across all ORFs. **B and C)** Redistribution of ribosome stalls across the B) *typA* or C) *ispH* open reading frames. Both genes show a clear drug-induced peak near the 5’ end and comparable overall translation efficiency. **D-G)** Codon-level analysis of stalling preferences within the top 1000 stall sites (D: TZD treated, F: DZD treated) or across the entire transcriptome (E: TZD treated, G: DZD treated). Individual stall sites were ranked as a function of log2 fold change in pause score between conditions, where pause score is defined as RPKM[codon]/ RPKM[gene]. **D, F)** pLOGO analysis utilized the top 1000 stall sites as foreground and the remaining sites as background. **E, G)** Transcriptome-wide heatmaps were produced by averaging the log2 fold change in reads per million for every instance of an amino acid occurring at positions –9 to +1 of a nascent chain.

Notably, no individual gene transcripts appeared to be particularly affected, with comparable translation efficiency (TE) scores, defined as a ratio between Ribo-seq and RNA-seq reads, in both treated and untreated samples. Despite relatively unchanged total TE scores (Extended Data Fig. 2A), clear drug-induced stalling peaks are visible, especially toward the 5’ regions of example genes *typA* and *ispH* (Fig. 2B,C).

We next analyzed bias in amino acid occurrence at the sites of TZD-mediated ribosome stalling. pLogo analysis of the top 1000 sites with increased ribosome occupancy in the presence of TZD revealed a clear dependence on the penultimate (−1) position of the nascent chain for stalling (Fig. 2D). In stark contrast to the well-documented Ala(−1)–dependent LZD-induced ribosome stalling preference^10,11,13^, we observed a significant preference for Ile in the –1 position (23.45% of the top 1000 sequences) in the presence of TZD. Besides Ile(−1), stalling was also observed when the penultimate position of the nascent chain in the drug-arrested ribosome was occupied by Gln (11.02%) and His (6.91%), indicating a dramatically distinct and also broader penultimate amino acid preference for TZD compared to LZD. As with linezolid, Gly is disfavored in the penultimate and C-terminal position of the nascent chain and as the A-site incoming amino acid.

To evaluate transcriptome-wide stalling preference of TZD, we averaged the change in frequency of occurrence of each amino acid within the ten C-terminal residues of the nascent chain and the incoming amino acid, across all stalling codons transcriptome-wide (Fig. 2E). In agreement with the trends observed with pLogo charts, the most pronounced changes in occurrence were observed for the nascent chain penultimate amino acid with a clear prevalence for Ile, His, or Gln in the TZD-arrested ribosome. To a lesser degree, stalling was also observed with Cys(−1), further emphasizing the broad –1 dependence of TZD-induced stalling. As with the top stall sites (Fig. 2D), we observed a strong preference against Gly(−1).

### Delpazolid exhibits enhanced stalling with nascent peptides containing penultimate Ile or His

Despite overall chemical similarity, our findings revealed distinct stalling preferences for TZD compared with LZD. A key distinguishing feature of the two antibiotics is the C5 moiety of the oxazolidinone ring, where the acetamidomethyl group of LZD is replaced by an alcohol moiety in TZD (Fig. 1A). We hypothesized that the nature of the C5 substituent may contribute to altered stalling preference. To test this hypothesis, we turned to ribosome profiling with DZD (Extended Data Fig. 3), an oxazolidinone compound active against methicillin-resistant *S. aureus* infections, as well as multidrug-resistant and extensively drug-resistant tuberculosis^29,33^. While the ring system of DZD is more comparable to that of LZD, DZD shares the truncated C5 hydroxymethyl moiety with TZD (Fig. 1A). DZD thus provides a valuable model system for exploring the role of the C5 substituent in determining context-specificity of oxazolidinones.

Much like TZD, DZD caused a buildup of ribosome occupancy early in ORFs (Fig. 2A). At the codon level, however, DZD-induced ribosome stalling sites showed a narrower preference in the nascent chain penultimate position compared to TZD. Analysis of both the top stall sites and transcriptome-wide stalling motifs resulting from DZD treatment showed a strong preference for Ile(−1) and His(−1) (33.03 % and 13.01% in the top 1000 sites, respectively; Fig. 2F,G). We also observed a preference for Val as the incoming amino acid and an exclusion of Gly, particularly at the –1 position.

Taken together, our findings from Ribo-seq experiments suggest that TZD and DZD preferentially stall translation after Ile or His have been incorporated into the penultimate position of the nascent peptide. This is in stark contrast to LZD and RZD, which strongly prefer Ala at this position, supporting the hypothesis that the chemical nature of the C5 side chain of the oxazolidinone dictates ribosome arrest preference.

### Context-dependency of TZD and DZD-induced translation arrest can be recapitulated in vitro

To validate the identified stalling motifs within specific sequence contexts, we used the ribosome profiling data to select five genes with representative ribosome stall sites where the nascent chain would contain either Ile or His in the penultimate position (Fig. 3A,C, Extended Data Fig. 4). While the effects of the two antibiotics are largely similar, DZD-induced peaks of increased ribosome occupancy appear to be generally more prominent and specific than TZD-induced stalls. These observations are consistent with a more narrow specificity of DZD-induced stalling (Fig. 2F,G), as compared to TZD (Fig. 2D,E).

**Figure 3:**
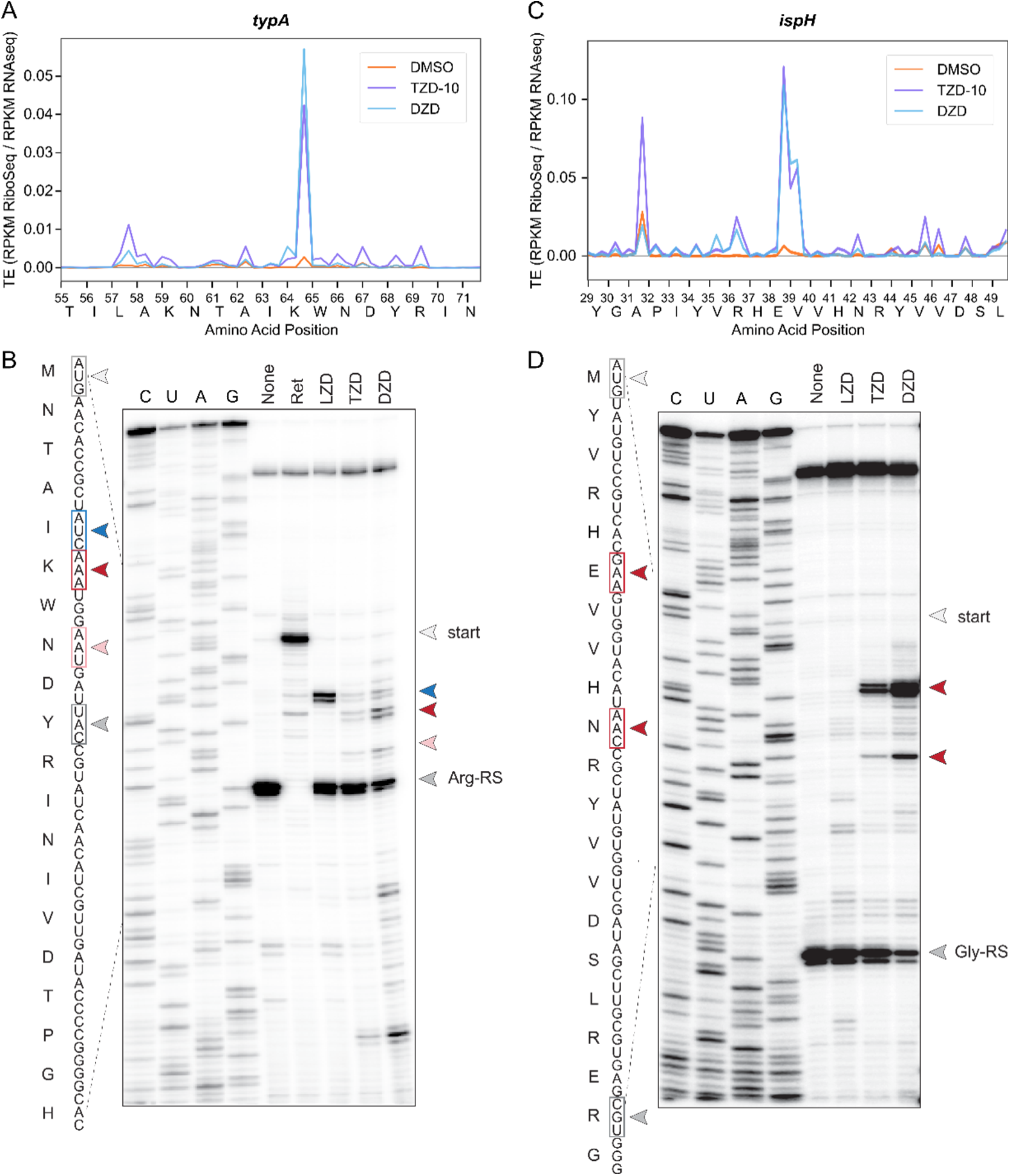
Validation of tedizolid and delpazolid-induced stall sites. **A)** Ribosome profiling traces of TZD (purple) and DZD (blue) indicate Ile(−1)-dependent stalling during translation of NTAIK sequence within the *typA* open reading frame compared to DMSO (orange). **B)** Ribosome toeprinting performed on the *typA*-derived NTAIK mRNA template in the presence or absence of LZD, TZD, or DZD. Stalling with Ala(−1) is denoted in blue, Ile(−1) in red, and Trp(−1) in pink. Pleuromutilin antibiotic retapamulin was used to induce stalling at the start codon (light gray). Arginine tRNA-synthetase inhibitor Arg-AMS (Arg-RS, dark gray) was included in all reactions to “catch” any ribosomes that progressed beyond the oxazolidinone-dependent stall sites. **C)** Ribosome profiling traces within the *ispH* ORF indicate TZD– and DZD-dependent stalling with His(−1) at YVRHEV site. **D)** Ribosome toeprinting performed on an *ispH*-derived mRNA template in the presence or absence of LZD, TZD, or DZD. Stalling with His(−1) is denoted in red. Glycine tRNA-synthetase inhibitor Gly-AMS (Gly-RS, dark gray) was included in all reactions to “catch” any ribosomes that progressed beyond the oxazolidinone-dependent stall sites. All compounds were treated at 50 μM for toeprinting reactions.

The selected stall sites were evaluated *in vitro* by toeprinting analysis. For these, we designed templates for in vitro translation that encoded 5-7 amino acids preceding the anticipated site of the drug-induced arrest, using either the gene’s natural start codon or introducing a new start codon at the appropriate distance from the stall site. After translation of the templates *in vitro* in the presence or absence of antibiotics, the sites of the ribosome arrest were determined by reverse transcriptase-mediated extension of the primer annealed 3’ to the arrest site.

The isoleucine-containing NTA**I**KW peptide derived from the *typA* gene showed a particularly strong drug-induced ribosome stalling signal in the Ribo-seq data and was therefore used in the initial toeprint experiments (Fig. 3A,B). In the presence of LZD, ribosome stalling occurred only when Ala4 of the template was in the penultimate position of the nascent peptide, consistent with the previously established stalling preference^10,11^. While we also observed a faint band at this position for TZD and DZD, both drugs also induced stalling at the next codon, when Ile appeared in position –1 of the nascent chain. Consistent with the ribosome profiling data, the Ile(−1) stalling bands were stronger under DZD treatment than under TZD treatment. An additional faint band placing Trp in the –1 position was observed (Fig. 3B). In agreement with the *in vivo* Ribo-seq data, ribosome arrest at the stall sites was incomplete, and a large fraction of ribosomes were able to reach the Arg11 “catch” codon, where they were arrested due to the presence of an Arg tRNA-synthetase inhibitor in the reactions.

To further evaluate oxazolidinone-induced Ile-1–dependent stalling, we carried out toeprinting analysis using the *dcuA* transcript, which contains the TZD– and DZD-induced stalling site within the VVELIIV motif (Extended Data Fig. 4A-C). In the cell-free translation system, two distinct stall sites were observed within this sequence in the presence of either TZD or DZD: one with penultimate Leu, and another with the first of two consecutive Ile residues in the –1 position. The exact placement of the drug-stalled ribosome within this template was further verified by using Ile tRNA-synthetase (Ile-RS) inhibitor mupirocin, whose presence causes ribosome stalling during decoding of A-site Ile. The penultimate Ile stall site induced by TZD and DZD is positioned one codon after the mupirocin-induced site, in line with their differing modes of action. These results confirm the importance of the penultimate Ile for stalling, while also suggesting a more relaxed sequence dependence of TZD and DZD action in the cell-free translation system in comparison with cellular translation.

We used Ribo-seq data to select two genes, *ettA* and *ispH*, to evaluate the importance of penultimate His for TZD– and DZD-induced stalling *in vitro* (Fig. 3C,D, Extended Data Fig. 4D-F). In contrast to LZD, DZD, and TZD treatment resulted in distinct toeprint bands within the *ettA* and *ispH* templates. These stalls placed the arrested ribosome one codon down from the respective His codons, thereby confirming that the penultimate His residue in the nascent peptide is conducive to the drug-induced translation arrest. These findings were further corroborated with the *rhlB* template (Extended Data Fig. 4G-I). As with the Ile(−1) stall sites, the arrest was more prominent with DZD than with TZD. Inspection of the stronger stall sites in each sequence revealed the presence of Arg preceding the penultimate His residue, matching an [R/K]HXX motif suggested by pairwise analysis of stall sites from the ribosome profiling for both TZD and DZD (Extended Data Fig. 5).

### Tedizolid stalling induces a novel stalled peptide conformation

Having shown that TZD and DZD exhibit Ile(−1) stalling, we aimed to identify the molecular basis of this altered specificity. The *typA*-derived NTA**I**K site was selected for structural analysis, as it represents one of the strongest stall sites identified by ribosome profiling. Stalled ribosome complexes (SRCs) were produced via *in vitro* transcription-translation of an MNTA**I**K template using *E. coli* ribosomes in the presence or absence of TZD. To favor the formation of a homogeneous complex and allow for the preparation of a drug-free ribosome-nascent chain (RNC) complex, the coding sequence was truncated after the Lys6 codon to generate a stop-less mRNA. The complex generated from the cell-free system was purified and applied to cryo-EM grids. Homogeneous refinement yielded a 2.16 Å resolution map of the TZD-MNTAIK SRC, with well-defined density for TZD (Fig. 4A,B, Table 1, Extended Data Fig. 6A).

**Figure 4:**
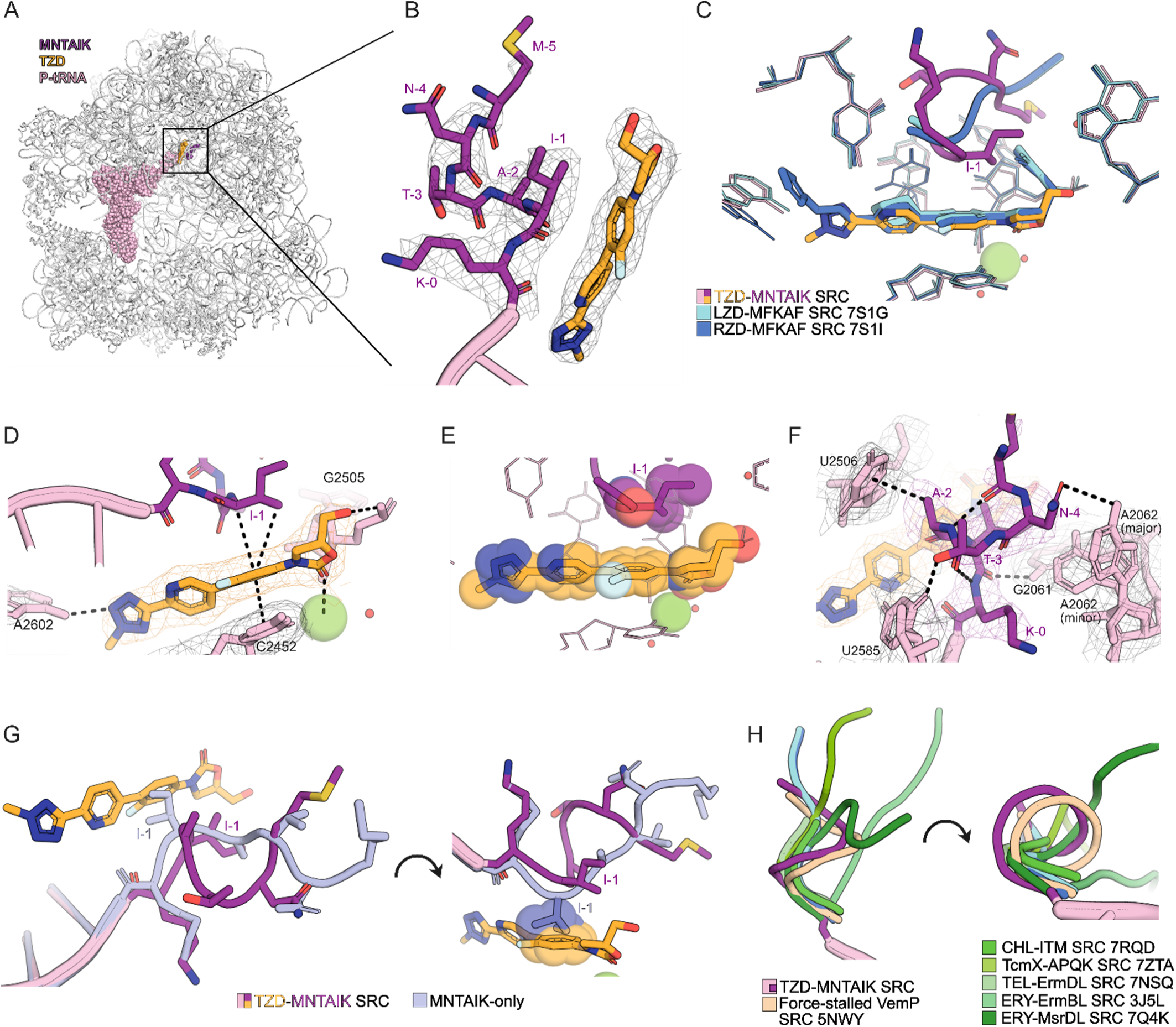
Tedizolid-induced ribosome stalling alters conformation of nascent peptide. **A)** Cross-section of the cryo-EM density map of the TZD-stalled 70S *E. coli* ribosome in complex with peptidyl-tRNA (TZD-MNTAIK SRC). **B)** Close-up of TZD (gold) in complex with MNTAIK arrest peptide (magenta). **C)** Alignment of TZD binding site and nascent chains with those of previous oxazolidinone stalled complexes: LZD-MFKAF SRC (PDB 7S1G, teal) and RZD-MFKAF SRC (PDB 7S1I, blue). **D)** Direct interactions between TZD and 23S rRNA (pink) or peptide (magenta). Dashed black lines represent binding interactions. **E)** Spheres representation of TZD interaction with penultimate Ile. Spheres contour highlights the shape complementarity between the drug and the peptide. **F)** Direct interactions between MNTAIK arrest peptide and 23S rRNA residues. **G)** Overlay of peptide path for the TZD-MNTAIK SRC and an analogous drug-free MNTAIK RNC (lavender). Right: spheres representation of the clash between TZD and Ile-1 of the drug-free peptide complex. **H)** Comparison of peptide path for TZD-MNTAIK SRC with other drug-stalled peptide complexes (green; PDB 7RQD, 7ZTA, 7NSQ, 3J5L, 7Q4K) and the force-sensitive stalling peptide VemP (beige; PDB 5NWY). Densities are shown at 3.0σ.

**Table 1:**
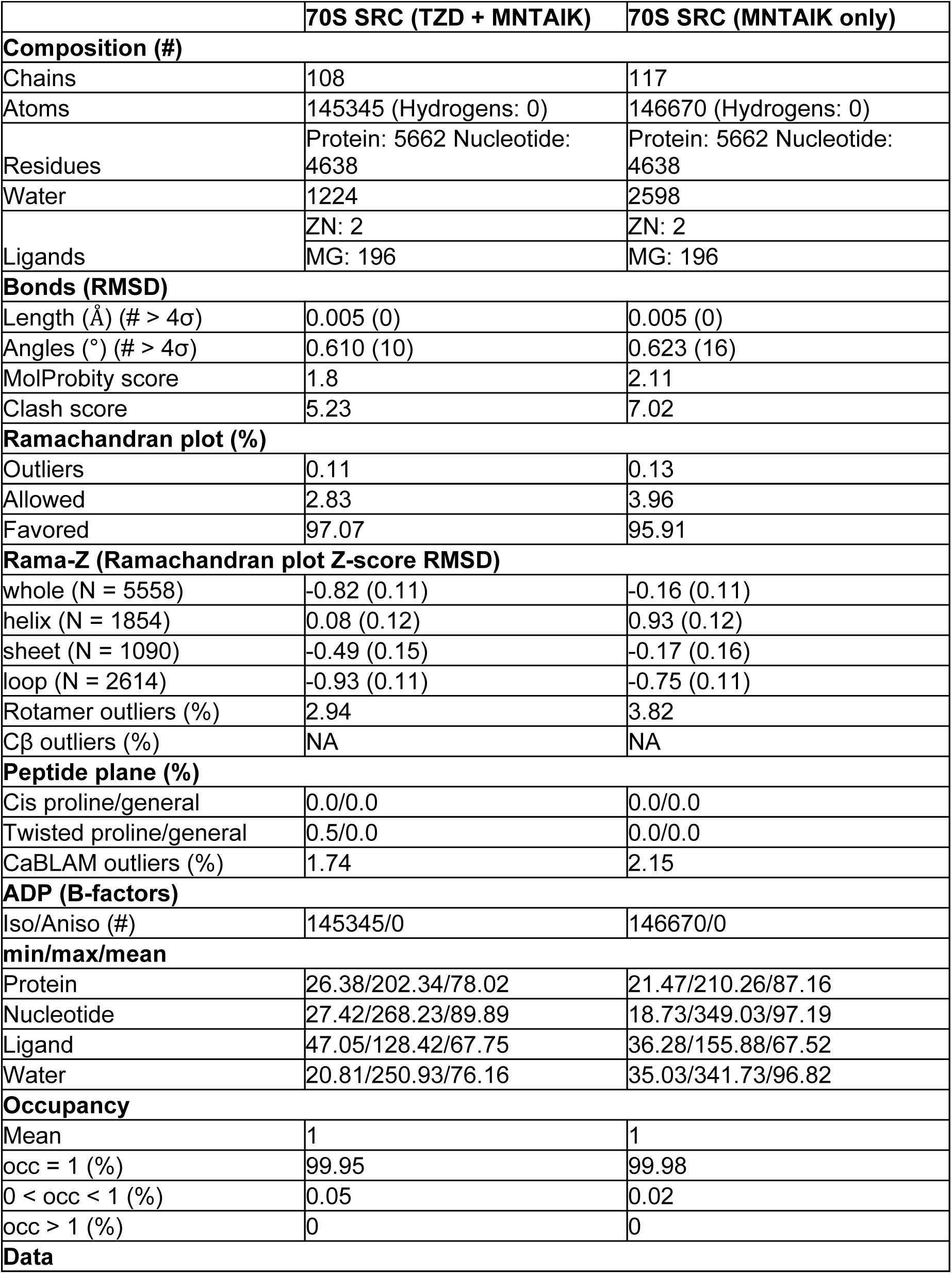
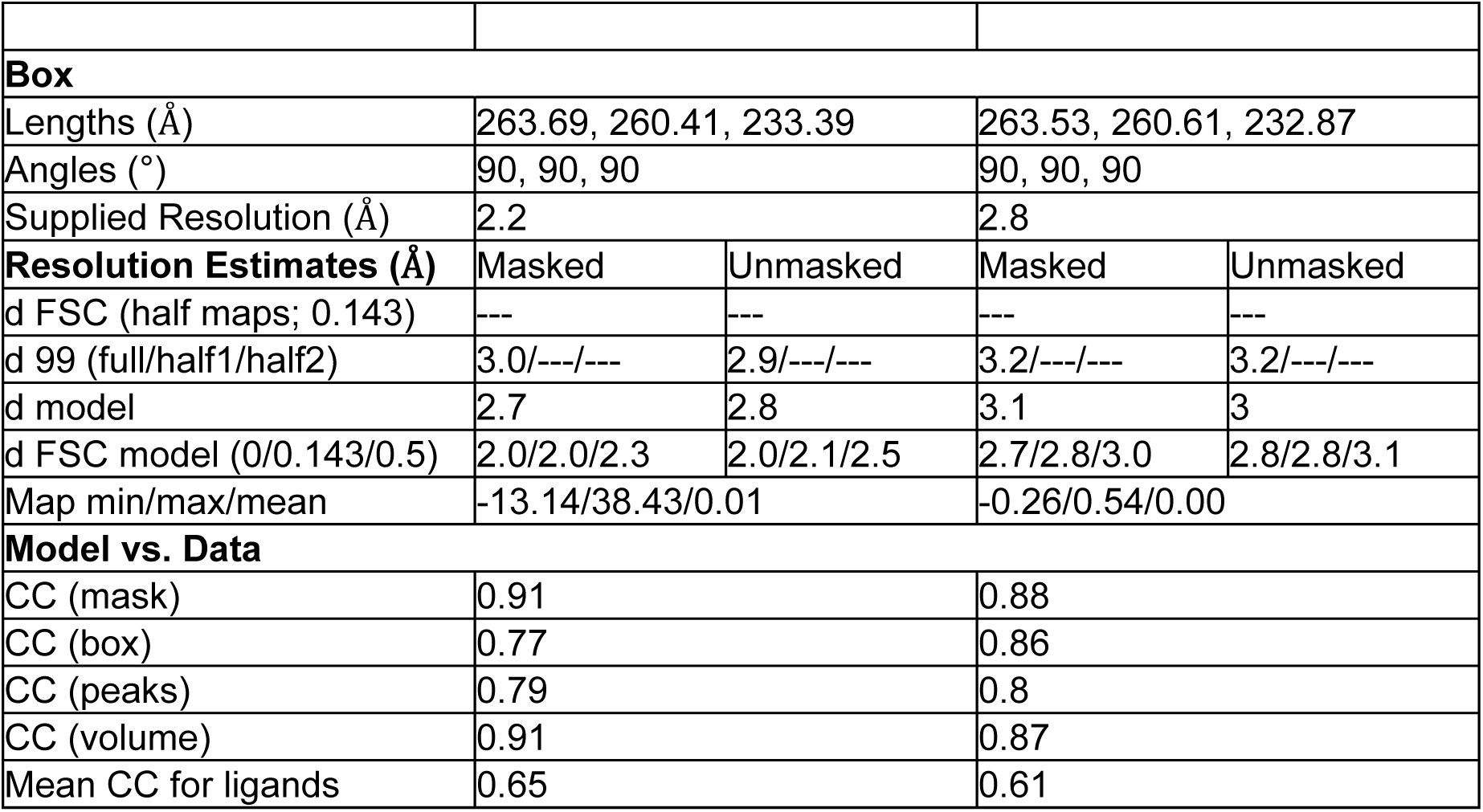
Cryo-EM Data Processing and Refinement.

While no density was observed for A– or E-site tRNA, the map contained clear density for the peptidyl-tRNA in the P-site. Modeling and real-space refinement unambiguously placed Ile in the penultimate position of the nascent MNTAIK peptide (Fig. 4B). This peptide assignment was further verified via analysis of density in the decoding center, which matched the AAA-Lys codon in the P-site of the decoding center (Extended Data Fig. 6C).

We next compared this TZD-stalled complex with previously published LZD-MFKAF and RZD-MFKAF SRC complexes. The A, B, and C rings of these drugs showed a near-identical placement (Fig. 4C). TZD binding is stabilized by a number of interactions with the ribosomal RNA in the stalled complex (Fig. 4D). In particular, π-π stacking between the oxazolidinone B-ring and C2452, as well as coordination of a nearby Mg^2+^ ion via the A-ring carbonyl moiety, are consistent with previously observed interactions in LZD and RZD^11^. The D-ring of TZD also forms a hydrogen-bond interaction with A2602. This residue is unavailable for interaction with the shorter LZD ring system and instead engages RZD via π– π stacking. However, we observed a major difference in the positioning of the C5 substituents emerging off the A ring. While the longer substituents of LZD and RZD fold toward the nascent peptide chain, the hydroxyl group of the shorter C5 hydroxymethyl substituent of TZD extends towards the exit tunnel wall, resulting in a wider peptide pocket (Fig. 4E) for TZD compared to LZD– or RZD-containing ribosome complexes. This rotation is supported by a hydrogen bond with the backbone of G2505 in 23S rRNA.

In contrast to the nearly identical positioning of the different oxazolidinones, the nascent chain in the TZD-SRC assumes a conformation that drastically differs from the largely extended conformation of the MFK**<u>A</u>**F nascent peptides in the LZD– and RZD-stalled complexes^11^ (Fig. 4C). The MNTA**I**K nascent peptide assumes an i+3 helical conformation, enforced by hydrogen bonding between the backbone amides of Lys(0) and Thr(−3) as well as Ala(−2) and Met(−5) (Fig. 4B,F). To test whether the unusual conformation is an intrinsic property of the MNTAIK sequence or whether it is specifically induced by the bound antibiotic, we trapped a drug-free complex with MNTAIK-peptidyl tRNA in the PTC using the analogous *in vitro* transcription-translation strategy in the absence of TZD (Table 1, Extended Data Fig. 6B). Although the resulting 2.77Å map of the drug-free complex showed poorer defined density in comparison to the TZD-MNTAIK complex, we were able to unambiguously assign and trace the peptide backbone (Extended Data Fig. 6B). In contrast to the helical peptide of the drug-stalled complex, we observe an extended nascent chain trajectory in the absence of TZD (13.7Å measured from Cα of Lys(0) to the Cα of Met(−5), versus 9.3Å in the TZD-stalled complex). Interestingly, this elongated conformation would place Ile(−1) in direct steric clash with the bound TZD (Fig. 4G, Extended Data Fig. 7). The comparison of the drug-free and drug-bound structures therefore illuminates an important concept – that PTC bound antibiotic is able to alter the trajectory of the nascent protein chain in the exit tunnel.

In the helical conformation stabilized by TZD, Ile(−1) can form two potential CH-π bonds with the B-ring of TZD via hydrogen atoms from its α or γ carbon (Fig. 4D). Distances and C-H_(pred)_-π angles for both of these putative interactions are consistent with established constraints for CH-π bonding^34^. We also observe a number of interactions between the nascent peptide and 23S rRNA, which may further facilitate this novel peptide conformation (Fig. 4F). In particular, the Ile(−1) backbone carbonyl forms a hydrogen bond with 23S nucleotide G2061, which has been noted to contribute to peptide positioning in the catalytically competent ribosome^1^. In addition, the side chain of Ala(−2) is within CH-π interaction distance from critical rRNA nucleotide U2506.

The path of the nascent peptide in this TZD-MNTAIK stalled ribosome complex appears to be unique amongst other drug-stalled complexes^9,11,12,18–20,22^ (Fig. 4H). Instead, the peptide path most closely resembles that of the force-sensitive stalling peptide VemP^35^. Collectively, these results indicate that drug binding can influence a peptide’s conformational state in the PTC, leading to translation stalling.

### Tedizolid-induced stalling results in reorientation of critical rRNA nucleotides

Reorientation of critical nucleotides in and around the PTC has been proposed to be at the heart of the mechanism of translation arrest resulting from cooperation between the antibiotic and the nascent peptide, although the involvement of various nucleotides differs by drug^11,18–20,22^. We thus compared the orientation of these nucleotides in our TZD-MNTAIK SRC versus the peptide-alone complex and an uninduced state of the ribosome, as well as the LZD-stalled ribosome complex (Fig. 5). Where the LZD SRC acts as a comparison for stalled conformation of the ribosome, our peptide-only structure represents a more catalytically competent state of the ribosome. A major reorientation is apparent in the functionally critical nucleotides U2585, A2062, A2602, and U2506.

**Figure 5:**
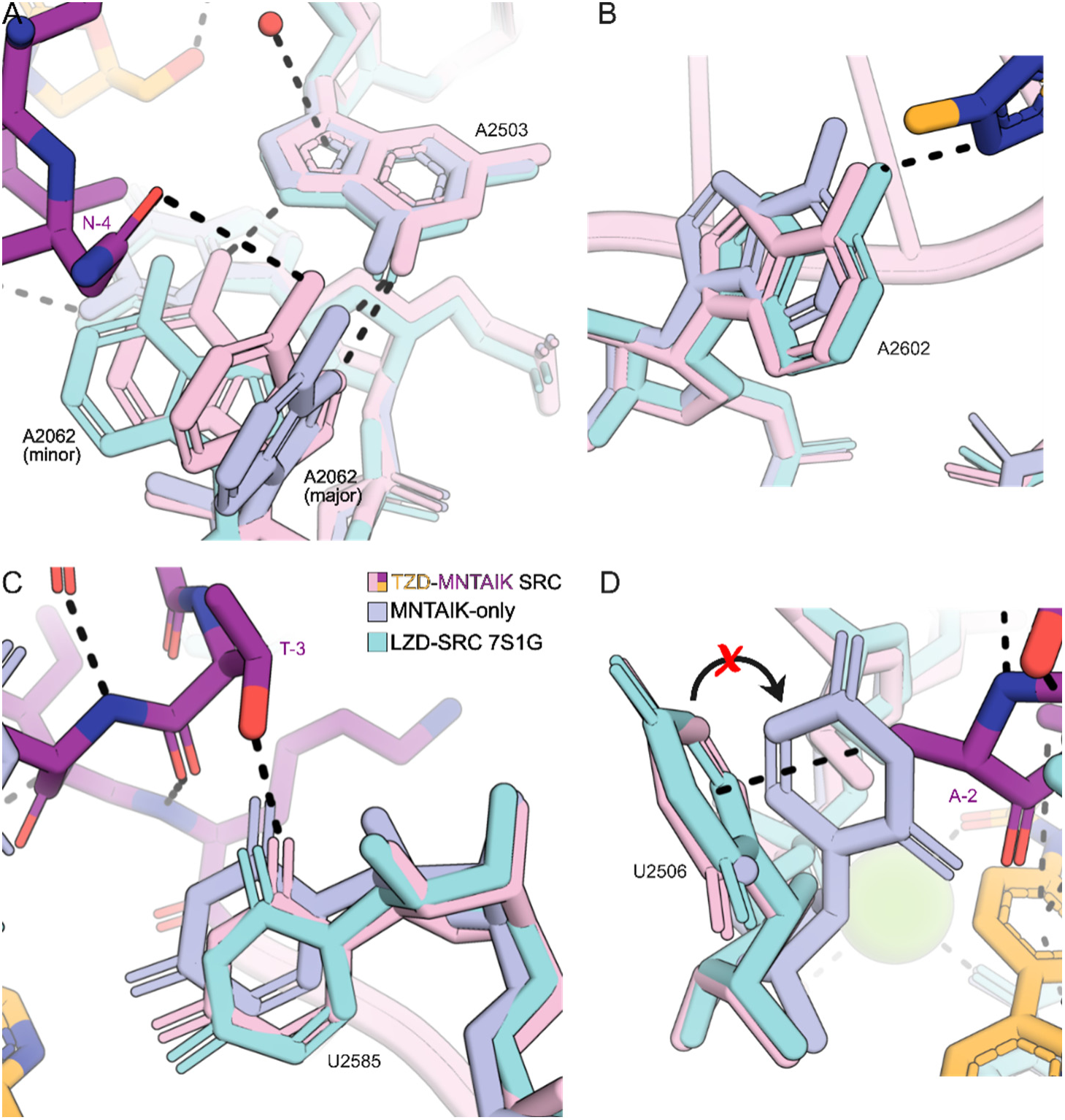
Ribosome stalling by tedizolid alters conformation of rRNA nucleotides. Comparison of 23S rRNA nucleotide positions in the TZD-MNTAIK SRC (rRNA, tRNA: pink, TZD: gold, MNTAIK peptide: magenta) versus the drug-free MNTAIK RNC (lavender) and previous LZD-MFKAF SRC (teal, PDB 7S1G). Putative interactions within the TZD-SRC are shown as black dashed lines. **A)** 23S rRNA nucleotide A2062 takes on two orientations in the TZD-MNTAIK SRC, mimicking either the MNTAIK-only or LZD-SRC conformation. **B)** A2602 shifts to accommodate a hydrogen bond with the D-ring of TZD, comparable to its position in the LZD-SRC. **C)** U2585 shifts toward the position observed in LZD-SRC to accommodate the nascent chain of T(−3). **D)** A predicted steric clash with Ala(−2) of MNTAIK peptide in the TZD-SRC prevents U2506 from accessing the internally-rotated conformation observed in the peptide-only complex.

In the majority of *E. coli* stalled complexes, the exit-tunnel nucleotide A2062 forms a non-canonical Hoogsteen base pair with the nearby nucleotide A2503. In the case of LZD– and RZD-SRCs, this paired conformation is further stabilized via a direct H-bond interaction with the drug^11^. The lack of direct interaction between A2062 and the TZD C5 substituent destabilizes this nucleotide, resulting in two conformational populations, akin to what has been previously observed in drug-only ribosome complexes^11^. We observe the paired conformation only as a minority population (40%) in the TZD-SRC. Instead, A2062 is rotated towards the tunnel lumen in its majority population (60%), a conformation that may be stabilized by a potential H-bond with the flexible Asn(−4) side chain of the nascent peptide (Fig. 5A, 4F). This rotation additionally precludes any interactions between A2062 and the peptide backbone previously observed in non-arrested complexes^1^. In comparison, we observe only the unpaired conformation of A2062 in the peptide-alone complex.

Drug-induced stabilization of the flexible PTC nucleotide A2602 has been shown to contribute to ribosome stalling by steric interference with incoming aminoacyl-tRNA^9,19,20,36^. Compared with the drug-free RNC, we observe a slight repositioning of the residue to accommodate TZD in the stalled complex (Fig. 5B). The newly adopted orientation closely resembles that of the LZD-SRC, despite the drug’s shorter ring system. Nucleotide U2585 also shifts in the TZD-SRC in comparison with the drug-free RNC to accommodate the novel peptide path, assuming a position similar to that observed in the LZD-SRC. In TZD-SRC, the newly adopted conformation may be stabilized by a possible hydrogen bond to the side-chain hydroxyl group of Thr(−3) in the nascent peptide (Fig. 5C, 4F).

Nucleotide U2506 is also significantly repositioned in the TZD-SRC relative to the drug-free RNC complex, resembling the positioning of this nucleotide in LZD-RSC. Stabilization of U2506 in this conformation is associated with a catalytically nonproductive state of the ribosome^20,37,38^. In the drug-free structure, we observe density for U2506 in the tunnel lumen. In this conformation, U2506 would clash significantly with the side chain of Ala-2 on the compacted nascent chain, resulting in the rotation of U2506 towards the tunnel wall in the TZD-SRC (Fig. 5D). By evading steric clash, this rotation of the nucleotide enables a CH-π interaction between U2506 and the side chain of Ala-2. The tunnel wall orientation of U2506 observed in the TZD-SRC is consistent with its orientation in the nonproductive state of the ribosome (Extended Data Fig. 8).

## DISCUSSION

Our investigation into the mechanism of translation inhibition by oxazolidinone antibiotics TZD and DZD shows that these antibiotics inhibit translation in a context-dependent manner, a feature previously observed for LZD and RZD. In contrast to a requirement for the presence of Ala in the penultimate position of the nascent chain for LZD and RZD-induced ribosome stalling, both TZD and DZD preferentially stall translation of nascent peptides containing Ile or His as penultimate residues. The stalling preference is extended for TZD to encompass Gln(−1) as well. The novel sequence preference of these oxazolidinones likely stems from the presence of a shorter C5 hydroxymethyl side chain in TZD and DZD, which allows for a wider peptide path and accommodation of larger side-chains at the penultimate amino acid residue of the nascent chain.

Structural analysis of the TZD-bound ribosome stalled with the Ile(−1) containing nascent peptide revealed a novel, compact trajectory of the growing protein chain, distinct from the extended conformations observed in other drug-stalled complexes analyzed to date^9,11,12,18–20,22^. This novel conformation represents the first instance of a drug-induced helical conformation of the nascent polypeptide immediately proximal to the P-tRNA. A similarly compact conformation containing a C-terminal helix in the nascent chain was previously observed in the force sensor protein VemP^35^. While the α-helical terminus of VemP is produced from a 20-amino acid stalling motif, TZD-induced stalling at the MNTAIK peptide achieves helical conformation of a far shorter peptide segment. This novel drug-induced conformation is particularly interesting in light of recent findings that ribosomes stabilize non-arrested peptides in an extended β-strand conformation for more efficient peptide bond formation^1^, suggesting that altered conformation is likely to play an important role in the mechanism of TZD-mediated inhibition of translation. The extended β-strand-like peptide trajectory observed for MNTAIK on the ribosome in the absence of TZD further supports this hypothesis.

The observed peptide trajectory appears to result from a combination of factors. The helical peptide conformation induced by the drug avoids steric clash between the peptide and TZD. This novel conformation, accommodated by an expanded nascent peptide-facing pocket in the drug, is stabilized by intramolecular hydrogen bonds and CH-π interactions between the Ile(−1) and TZD. We previously showed that the acetamidomethyl group of LZD sterically occludes a portion of the PTC, creating a smaller binding pocket that can accommodate the side chain of Ala^11^. In contrast, the C5 hydroxymethyl substituent of TZD is pulled towards the exit tunnel wall, thereby widening the amino acid side chain binding pocket. Locked into this conformation by an H-bond interaction with the rRNA backbone, ribosome-bound TZD is less restrictive of larger penultimate residues in the nascent peptide. The compacted nascent peptide path precludes formation of a H-bond network previously identified as critical to proper positioning of the peptide for catalysis of peptide bond formation^1^; structures of other stalled complexes with alternate or disrupted C-terminal secondary structure^35,39–41^ suggest that disruptions to this network may contribute to stalling^1^. Further structural studies will be necessary to evaluate TZD-induced stalling of His(−1)-containing nascent chains, though we hypothesize that accommodation of this larger residue will also benefit from a relatively widened exit tunnel.

While both TZD and DZD show a preference for Ile(−1) or His(−1) at their stall sites, we noted critical differences between the stalling preferences of the two drugs. TZD-mediated stalling is inclusive of a wider variety of alternative penultimate nascent chain residues compared to DZD, including a similarly intense preference for Gln(−1) and weaker stalling with Cys(−1). This difference is likely attributed to the presence of the D-ring in TZD, which, through an added interaction with the ribosome, may enhance stalling even at less favorable stall sites. In addition to the narrower –1 stalling preference, DZD-induced stalling peaks tend to be higher and less frequent than those caused by TZD. This trend was recapitulated by the *in vitro* toeprinting experiments. As a result, treatment with DZD leads to a greater fraction of ribosomes stalling at fewer sites, rather than being distributed across many.

Not every occurrence of Ile, His, or Gln in the nascent peptide leads to pronounced TZD-or DZD-induced translation arrest (Extended Data Fig. 9), suggesting that a longer sequence context may significantly contribute to stalling, as previously observed for other antibiotics^15,20^. Although TZD and DZD appear to induce stalling at the majority of Ile(−1) and His(−1) sites tested by toeprinting, we noted variation in band intensities for sites containing multiple potential stalling motifs (Fig. 3B,D, Extended Data Fig. 4C,F). Pairwise analysis of the ribosome profiling datasets provided new insights, revealing an [R/K]HXX motif at the stronger His(−1)–containing stall sites of *ispH* and *ettA* (Fig. 4D, Supplementary Fig. 4F, 5A-B). This motif, shared between the two C5-hydroxymethyl oxazolidinones, represents the first instance of expanded specificity beyond the –1 position for this class of compounds. Further exploration of the molecular basis of this pairwise preference may reveal why certain instances of the stalling motif are favored over others.

TZD stalled complex revealed unique interactions of this antibiotic with the ribosome. The C5-hydroxymethyl group of the drug forms a hydrogen bond with G2505. Although the resulting orientation of the C5 substituent towards the tunnel wall eliminates any direct drug interaction with A2062, the minor conformation of this base engages in non-canonical base pairing with A2503. Interestingly, the major conformation places A2062 near the tunnel lumen, where it is capable of supporting the compacted peptide conformation via a putative interaction with Asn(−4) side chain. At the distal end of the oxazolidinone, nucleotide A2602 forms an H-bond with TZD, which may aid in stabilizing antibiotic binding to the ribosome. The observed interaction highlights the versatility of A2602 in stabilizing antibiotic binding, as the same rRNA nucleotide stabilizes radezolid binding via a π-π interaction with its D-ring^11^. In addition, stabilization of this A-site–oriented nucleotide position likely interferes with proper accommodation of incoming aa-tRNA, further contributing to inhibition of translation^9,19,20,36^. We also observed reorientation of nucleotides U2506 and U2585, despite a lack of direct interaction with the drug. The resulting conformations closely resemble those of catalytically incompetent ribosome complexes^22,35,37,42,43^, including that of LZD-SRC^11^, providing further insights into the mechanism of stall induction by TZD.

We have shown that changes in the structure of oxazolidinone antibiotics can lead to major alterations in stalling sequence preferences. In addition, our findings reveal a surprising drug-induced malleability of the peptide path traversing the PTC. PTC-bound oxazolidinone modulates peptide path, thereby contributing to allosteric reorientation of critical rRNA residues, preventing continued translation. An exploration of context-specific stalling using an expanded library of oxazolidinones will elucidate how defined structural alterations affect the specificity of context-specific ribosome stalling and the trajectory of the nascent peptide.

Given the importance of context-specific ribosome stalling in mediating the expression of antibiotic resistance factors^18–20,22,44,45^, a better understanding of the molecular contributors to this mechanism is critical to developing antimicrobial compounds capable of averting the induction of resistance genes. In particular, the shared Ala(−1) specificity of LZD and the natural product chloramphenicol (CHL)^10^ makes LZD likely to activate expression of CHL-inducible resistance genes^45,46^. With altered stalling specificity, TZD and DZD may prevent this activation and thus maintain their antimicrobial activity against a wider range of otherwise resistant bacterial strains. Because activation of many resistance genes relies on the amino acid specificity of the inducing antibiotic, modifications to such specificity could be implemented to improve therapies for multidrug-resistant bacterial infections.

## METHODS

### Antibiotics

Oxazolidinones were purchased from Ambeed (tedizolid A139024, delpazolid A194738, linezolid A360550).

### Ribo-Seq & parallel RNA-seq

Ribosome profiling was performed as described previously^16,47^, with minor modifications.

Primer sequences are listed in Supplementary Table 1. *E. coli* BW25113 *ΔacrB* were grown overnight at 200 rpm & 37°C in MOPS EZ Rich media (Teknova M2105). Cultures were diluted 1:200 and grown to OD600 ∼ 0.3-0.5. Oxazolidinones, dissolved in DMSO at 50x final concentration, were added to log-phase culture at a final concentration of 10x MIC (20 μg/mL TZD, 80 μg/mL DZD). Equal amounts of DMSO were added to the ‘no drug’ control cultures (2% DMSO final concentration). Cultures were shaken for 2.5 min, then filtered through a 0.2 µM nitrocellulose membrane (Whatman 7182-009), and the filtrate was rapidly flash-frozen. Cells were mechanically lysed on Qiagen TissueLyser II with 650 µL frozen lysis buffer [20 mM Tris pH 8.0, 10 mM MgCl2, 100 mM NH4Cl, 5 mM CaCl2, 0.4% TritonX-100, 0.1% NP-40, 100 U/mL RNase-free DNase I (Roche 04716728001), 10 U/mL SUPERase-In RNase Inhibitor (Invitrogen AM2696), antibiotic 10x MIC]. Canisters were pre-chilled in LN2, and lysis was performed over 5 cycles of 3 min at 15 Hz. Pulverized lysate was thawed and clarified by centrifugation at 20,000 xg for 10 min at 4°C. 12.5 AU clarified lysate (∼0.5 mg RNA) was digested for 1 hr at 25°C and 1400 rpm with 750 U MNase (Roche 10107921001), and the reaction was quenched with 5 mM EGTA and placed on ice. Clarified lysate was also aliquoted for RNA-seq. Digested samples were then applied to a 10-50% sucrose gradient (20 mM Tris pH 8.0, 10 mM MgCl_2_, 100 mM NH_4_Cl, 2 mM DTT, 10 U/mL SUPERase-In) and centrifuged at 201,000 xg for 2.5 hr at 4°C. Polysome profiles were obtained using the BioComp Fractionator, and monosome-containing fractions were collected and pooled. Combined monosome samples were purified by hot phenol extraction and 20 µg of sample was resolved by 15% TBE Urea gel (Novex EC6885BOX). Ribosome-protected mRNA fragments 15-45 nt in length (guided by primers SC-15 & SC-45) were excised and eluted from gel (RNA Elution buffer: 300mM NaOAc pH 5.5, 1mM EDTA pH 8.0) for library preparation as described below.

For RNA-seq performed in parallel, an additional 12.5 AU clarified lysate aliquots were thawed and hot-phenol extracted. Small RNA and tRNA were removed from the total RNA via MegaClear Transcription Clean-Up kit (Invitrogen, AM1908), and rRNA was depleted using MicrobExpress Bacterial mRNA Enrichment kit (Ambion, AM1905). Remaining mRNA was fragmented using RNA Fragmentation Reagents (Ambion, AM8740) under stringent conditions (95°C 1 min 45 sec). Fragments 20-100 nt in length were selected and purified from 20 µg sample on 15% TBE Urea gel (Novex EC6885BOX) with RNA Elution buffer and applied to the library preparation pipeline below.

#### Library preparation

Total size-selected mRNA was dephosphorylated using 10 U T4 polynucleotide kinase (NEB M0201L) at 37°C for 1 h. Dephosphorylated RNA was ligated to 0.4 µL of 50 µM Universal miRNA cloning linker (NEB S1315S) using 200 U of T4 RNA ligase 2 truncated KQ (NEB M0373L) for 3 h at 37°C. Ligated products of appropriate size range (32-62 nt) were isolated from 15% TBE Urea gel (Novex EC6885BOX) and eluted with RNA Elution buffer. RNA was converted to cDNA by reverse transcription with 200 U SuperScript IV RT (ThermoFisher 180900510) and 1.25 µL of 10 µM primer Ni-Ni-9. RT products were resolved on 10% TBE Urea gel (Novex EC6875BOX) and the appropriate size bands (111-141 nt) were eluted out (DNA Elution buffer: 10 mM Tris pH 8.0, 300 mM NaCl, 1 mM EDTA pH 8.0). Circularization of cDNA was performed using 100 U CircLigase ssDNA ligase (Epicentre CL4115K) at 60°C for 1 h. Ribosomal RNA was removed from ribosome profiling samples by incubating with biotinylated primers b-rRNA-1 through b-rRNA-5 and pulling down with MyOne Streptavidin C1 Dynabeads (Invitrogen 65001) to remove corresponding cDNA. Remaining cDNA was PCR amplified with primer P5 and barcoded P7_index primers to add on NGS handles for 5-9 cycles. Amplified DNA (154-184 bp) was then purified from 10% TBE gel (Novex EC6275BOX) and eluted in DNA elution buffer. NGS-amplified products were evaluated on Agilent Tapestation with D1000 kit (ScreenTape 5067-5582, reagents 5067-5583, ladder 5067-5586), pooled, and submitted for sequencing at the UCSF CAT on NovaSeqX with 100 cycle single end sequencing.

### Profiling Data Analysis

All profiling data processing and analysis was performed using custom scripts and referencing previous analysis methods^15,16,30^.

#### Preprocessing

Raw, demultiplexed reads from next-generation sequencing were trimmed to exclude 3’ linkers (CTGTAGGCACCATCAAT) using cutadapt version 4.4^48^. Reads with fewer than 15 or greater than 45 nt were excluded from downstream analysis. Sequences of tRNA, rRNA, and tmRNA were copied from the parent strain genome (CP009273.1^49^) to a new fasta file along with ladder (SC-15 and SC-45) and illumina (P5, P7_iXX) primer sequences. This collated file was used to exclude the described sequences from downstream analysis via Bowtie^50^ alignment with a maximum of 2 errors per alignment. Remaining sequences were then Bowtie aligned to the full parent strain *E. coli* BW25113 genome with a maximum allowable error of 2 nt and a maximum of one alignment location within the genome. Bowtie output files were converted to Wig format by aligning reads to their 3’ ends, trimming to remove end mismatches and filtering to exclude resulting reads less than 15 nt long. FastQC^51^ was performed at each stage to verify quality.

#### Metagene Analysis

P-site assignment and periodicity analysis were performed essentially as described previously^15^. Periodicity was measured by averaging reads at every third position within coding sequences (CDS) transcriptome-wide, excluding the first 18 nt of each CDS, and comparing between each of the three nt positions. Low-expression and coverage genes (average reads/position ≤ 0.5 and < 20% nonzero reads) were excluded from this analysis. Genes were further filtered to exclude CDS < 100 nt long for the following metagene analyses. Reads were analyzed around the start and stop codons of open reading frames, and the P-site was assigned at –15 nt from the 3’ end of measured reads. Read distribution analysis across ORFs was performed as described previously^32^. Nucleotide positions within genes were divided by full gene length and reads were assigned to the resulting fractional position (rounded to two decimal places). Reads were averaged across all ORFs under analysis and plotted.

#### Gene-level Analysis

P-site aligned reads with an offset of –15 nt were used for all analyses, and gene-level analyses were performed essentially as described previously^15^. Gene RPKM (reads per kilobase per million) scores were calculated by summing reads over the full ORF for and normalizing to gene length and total sequencing depth of the sample. Ribosome profiling RPKM scores were further normalized to RNA-seq RPKM scores, resulting in a measure of translation efficiency (TE) which was used for correlation analysis. Where indicated, TE for each gene was averaged across two replicates for each experimental condition. Statistical significance of differences in TE for each condition compared to DMSO control was calculated using independent t-tests for each gene with Benjamini-Hochberg FDR correction. Data was filtered to exclude ORFs < 18 nt, with <20% nonzero reads, and ≤ 0.5 average reads/position, and genes not present in both conditions being analyzed after application of these filters were eliminated from analysis.

#### Codon-level analysis

Codon analyses were performed following the same filters as the gene-level analyses. Further filters were applied to exclude the terminal 2-10 codons of each CDS and codons with fewer than 5 reads total. Codons not appearing in both conditions being analyzed were removed from consideration. Codon reads were normalized to total gene reads to produce pause scores for each codon, and pause scores were averaged across replicates. Average pause scores were sorted on the basis of log2 fold change compared to DMSO.

pLOGO analysis^52^ was performed using the top 1000 stalling codons as foreground and all remaining codons as background, selecting an 11 amino acid window with the P-site codon peak at position 10 (−9:+1). For transcriptome-wide heat maps, codon RPM (reads per million) was calculated by normalizing codon reads to total sequencing depth. These codon RPM scores were applied at each amino acid in an 11mer window (−9:+1) around the peak, and individual amino acid frequency was calculated at each position by averaging RPM scores across all codons and replicates. Heat maps were plotted as the log2 fold change in averaged RPM scores compared to DMSO control. Pairwise heatmaps were produced to evaluate the frequency of amino acid pairs at the –2:-1 positions. For each potential amino acid pairing, codon RPM scores were averaged across all codons and replicates, and the log2 fold change in averaged pairwise RPM was calculated compared to the DMSO control.

### Toeprinting

DNA templates used in the toeprinting experiments were prepared by PCR using AccuPrime DNA Polymerase (Thermo Fisher). The sequences of the primers used are listed in Supplementary Table 1. The *ettA, ispH, typA, dcuA, and rhlB* templates were amplified from *E. coli* genomic DNA using universal primer T7-IR-fwd, and gene-specific primers IR-xxxX and NV1-xxxX-rev, where ‘xxxX’ represents the name of the corresponding gene. The artificial MFKAFKNIIRTRTL template was generated by a 5-primer PCR using primers T7, NV1, FWD, FKAFK and FKAFK-Rev as described^13^.

Toeprinting reactions were carried out in 5 µL of PURExpress transcription-translation system (New England Biolabs) essentially as described previously^53^. Antibiotics and aminoacyl-synthetase (RS)-inhibitors (mupirocin for Ile-RS or Arg-AMS and Gly-AMS for Arg-RS and Gly-RS, respectively) were added to the final concentration of 50 µM unless otherwise indicated. Following 30 min of translation, reverse transcription was carried out for 10 min using toeprinting primer NV1. cDNA products were separated on 6% sequencing gels. The gels were dried, exposed to a phosphorimager screen, and visualized in a Typhoon phosphorimager (GE Healthcare).

### GFP Translation Inhibition

*In vitro* transcription-translation reactions were carried out in 5 μL of the PURExpress system (NEB) containing 10 ng/μL of template DNA encoding the fluorescent protein mEGFP (gifted from the Seiple Lab) and supplemented with SUPERase-In RNase Inhibitor (Invitrogen AM2696) and the antibiotic of interest. Reactions were prepared on ice as a master mix of PURExpress and SUPERase-In which was added to a clear 96-well PCR plate (Bio-Rad MLL9601) containing plasmid and antibiotic or DMSO. Plates were sealed with clear covers (Bio-Rad MSB1001) and incubated at 37°C on CFX Duet Real-time PCR System (Bio-Rad) with fluorescent readings recorded every 5 min for 3 hr in the FAM channel (λ_ex_=450–490 nm, λ_em_=515–530 nm).

### Ribosome Purification

*E. coli* MRE600 were cultured overnight in LB media at 37°C and 220 rpm, then diluted 1:150 in LB and grown to OD ∼0.4-0.5. Cells were harvested by centrifugation at 4°C 4000 rpm for 15 min. Pellets were placed on ice and washed briefly (20 mM Tris pH 7.5, 100 mM NaCl, 10 mM MgCl2), divided in half, and flash frozen. Pellets were gently resuspended on ice in 15 mL Buffer AE (20 mM Hepes-KOH (pH 7.5), 200 mM NH_4_Cl, 20 mM Mg(OAc)_2_, 0.1 mM EDTA, 6 mM β-mercaptoethanol, 10 U/mL SUPERase-In) + 0.1 mM PMSF and filtered through a cell strainer. Filtered cells were lysed by 4 passages through an LM10 Microfluidizer (Microfluidics) at 10,000 psi. PMSF, BME, and SUPERase-In were supplemented to account for added volume. Lysate was incubated with 5 U/mL RQ1 RNase-free DNase (Promega M6101) for 15 min at 4°C, then spun down at 30,000 xg for 30 min and S30 fraction supernatant was collected. Tight-coupled ribosomes were purified from S30 fraction via 4x 32% sucrose cushion prepared in buffer 1E (20 mM Hepes-KOH (pH 7.5), 500 mM NH_4_Cl, 20 mM Mg(OAc)_2_, 0.5 mM EDTA, 6 mM β-mercaptoethanol, 10 U/mL SUPERase-In) via 16-18 hr centrifugation at 100,000 xg (28,000 rpm) in Beckman SW41Ti rotor. After removing supernatant, pellets were recombined by resuspension in 1 mL total buffer AE over ∼4 hr with slow rocking at 4°C. Remaining non-resuspended particles were cleared by centrifugation at 4°C for 10 min at 10,000 xg and ∼400 pmol loaded onto each of 6x 15-30% sucrose gradients prepared in buffer AE. Gradients were spun at 75,416 xg (21,000 rpm) for 16-18 hr at 4°C in a Beckman SW41Ti rotor. Fractionation was performed using BioComp Gradient Fractionator and monosome fractions were collected and combined. Ribosomes were precipitated at 4°C by slow addition of 30% PEG 20,000 in buffer AE to reach 9% PEG final concentration, then centrifuged at 17,500 xg for 10 min at 4°C to remove supernatant. Ribosomes were resuspended in buffer C (50 mM Hepes-KOH pH 7.5, 150 mM KOAc, 20 mM Mg(OAc)_2_, 7 mM BME, 20 U/mL SUPERase-In) to ∼13.3-20 pmol/μL by rotating at 4°C overnight and 5 μL aliquots were flash frozen in liquid nitrogen for *in vitro* transcription/translation reactions and structural studies.

### Stalled Complex Preparation

Non-stop templates for stalled complex preparation are composed of a T7 promoter for use with NEB PURExpress *in vitro* transcription/translation system, a Shine-Dalgarno sequence, and an ORF sequence containing the putative stall site but lacking a stop codon. Templates were synthesized by 4-primer PCR by combining equal volume of a 100 μM SRC-fwd-short, 10uM SRC-fwd-long, 100uM SRC-rev-short, and 10uM SRC-rev-long for the appropriate ORF sequence (Supplemental Table 1), along with 10mM dNTPs (NEB), and amplifying 30 cycles with T_a_ = 52°C using Phusion High-Fidelity DNA Polymerase (NEB M0530L). PCR products were purified using MinElute PCR Purification kit (Qiagen 28006).

Stalled complexes were formed for cryo-EM structural studies by *in vitro* transcription/translation. 100 μL reactions composed of PURExpress Δribosome kit (NEB E3313S) with 0.8 U/uL SUPERase-In, 100 ng DNA template, 240 pmol purified MRE600 ribosomes, and 5000 pmol (20x molar excess) oxazolidinone or DMSO to 0.5% were prepared on ice and incubated at 37°C for 1 hr. The reaction was halted by moving to ice, then diluted to 190 μL in ice cold Buffer C and loaded onto a 10-50% sucrose gradient prepared in buffer C. Following centrifugation in a Beckman SW41Ti rotor at 22,000 rpm for 16 hr at 4°C, the gradient was divided into 20 fractions using the BioComp Gradient Fractionator and monosome fractions were collected and combined. Ribosomes were precipitated with PEG 20,000 by slow addition at 4°C to a final concentration of 8% PEG, then spun down at 17,500 xg for 10 min at 4°C. The pellet was gently washed with 500 μL buffer C to remove residual sucrose, then resuspended in buffer C to a final concentration of ∼1-3 μM by rotating overnight at 4°C. Resuspended complexes were incubated 1 hr at 4°C with 30x molar excess of the appropriate antibiotic in a final concentration of 0.5% DMSO. This incubation step was omitted for the peptide-only complex. Prior to freezing grids, complexes were filtered through 0.22 μm PVDF filter (Millipore UFC30GV00) by centrifugation at 14,000 xg for 5 min.

### Cryo-EM

#### Cryo-EM sample preparation and data acquisition

After concentrating the sample to 400 nM, it was blotted on a Quantifoil R1.2/1.3 Cu grid with 2 nm continuous carbon. Blotting was carried out on FEI vitrobot Mark IV at 90% humidity with a blot time of 8s. Cryo-EM data was collected on a Titan Krios transmission electron microscope at 300 kV using a nine-hole beam image-shift scheme with coma compensation and automated acquisition via SerialEM (v4.1)^54–56^.

#### Cryo-EM image processing

Image processing involved binning of the movies by a factor of two, motion correction with MotionCor2^57^. Contrast transfer function (CTF) estimation was carried out using CTFFIND4^58^ and particles were picked using the template-based approach in CryoSPARC. We used PDB ID 7K00 as a starting structure to generate reference volume which was followed by 2D classification^59^. The particle box size was selected using previously established criteria^60^. Homogeneous refinement was used with per-particle CTF and defocus refinement after discarding the poor particles after 2D classification^61^.

#### Atomic model building and refinement

The initial atomic coordinates, derived from PDB entry 7K00, were docked into the cryo-EM density map via rigid-body fitting using UCSF ChimeraX. To facilitate the accurate modeling of non-protein moieties, ligand stereochemical restraints were generated through the eLBOW module^62^ within the Phenix software suite. Structural optimization was achieved through a cyclical process of manual rebuilding and local real-space fitting in Coot^63^, followed by automated global refinement against the experimental density using phenix.real_space_refine^64^. To ensure high-fidelity ligand geometry and improve the accuracy of electrostatic interactions, this refinement was further supplemented with force-field-based calculations employing the OPLS3e/VSGB2.1 model^65^. Two rounds of focused 3D classification were carried out using a mask for the tRNA and the peptide followed by a final round of homogeneous refinement. Final structural visualizations and figure rendering were executed in PyMOL (v3.0.2)^66^ utilizing unsharpened maps for the final density representations.

## EXTENDED DATA

**Extended Data Fig. 1.**
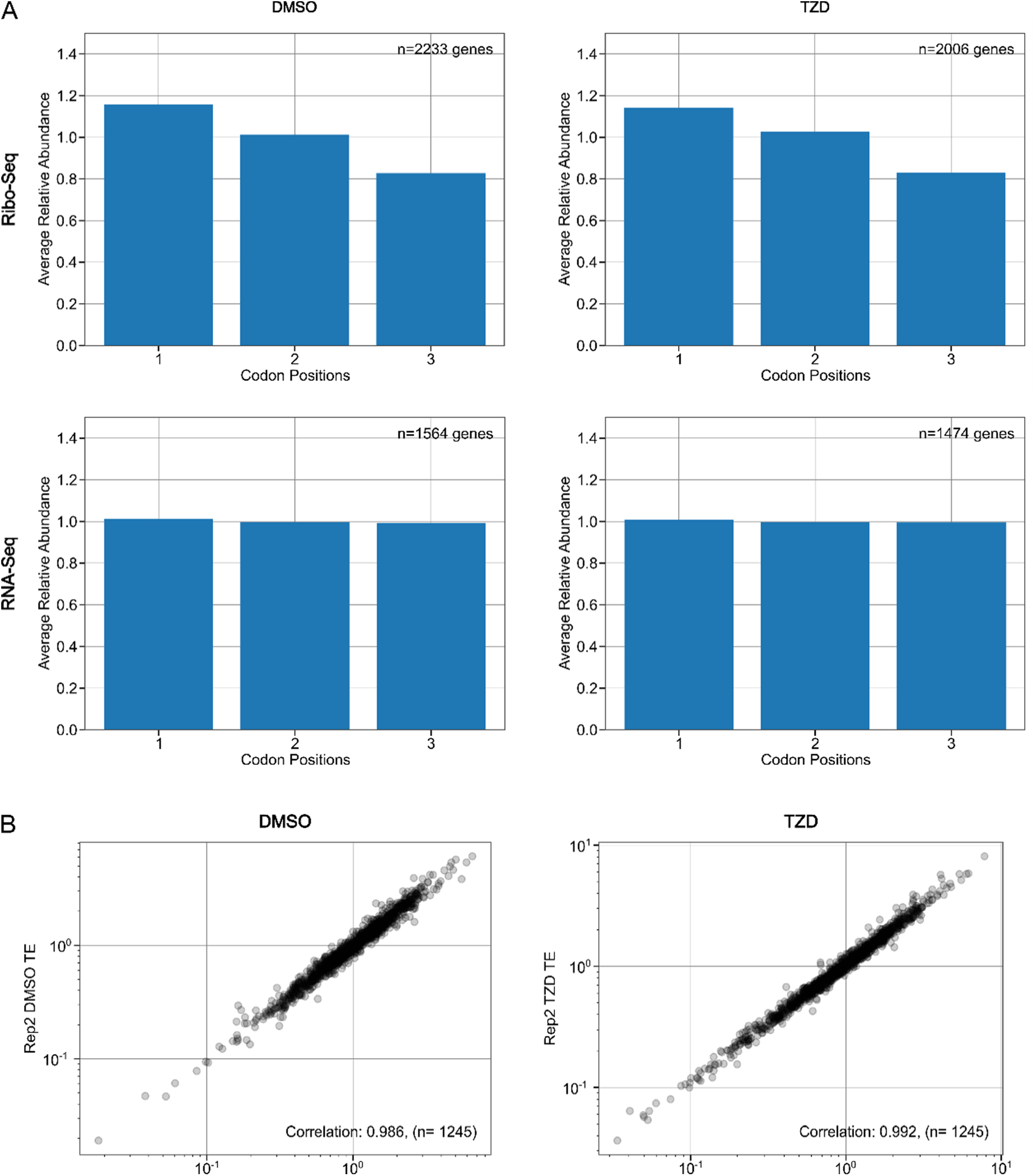
Ribosome Profiling Processing After TZD Treatment. **A)** Periodicity analysis of profiling data following 2.5 min treatment with DMSO or 10x MIC TZD. Top: Ribosome profiling. Bottom: RNA-seq. **B)** Genewise correlation between replicates. Changes in translation efficiency (RPKM Ribosome profiling/RPKM RNA-seq) for individual genes between experimental replicates following DMSO (left) or TZD (right) treatment.

**Extended Data Fig. 2.**
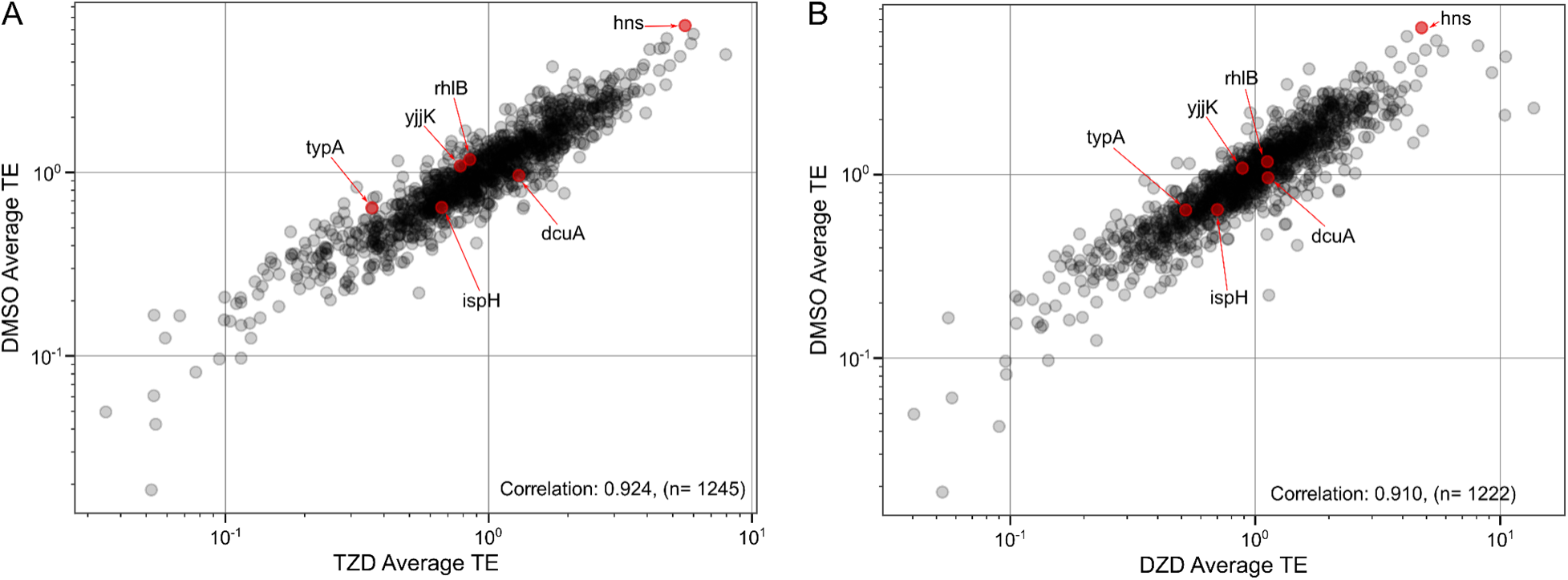
Genewise correlation in translation efficiency between. **A)** TZD-treated or **B)** DZD-treated conditions and DMSO control via ribosome profiling. Genes selected for toeprinting analysis are highlighted in red.

**Extended Data Fig. 3.**
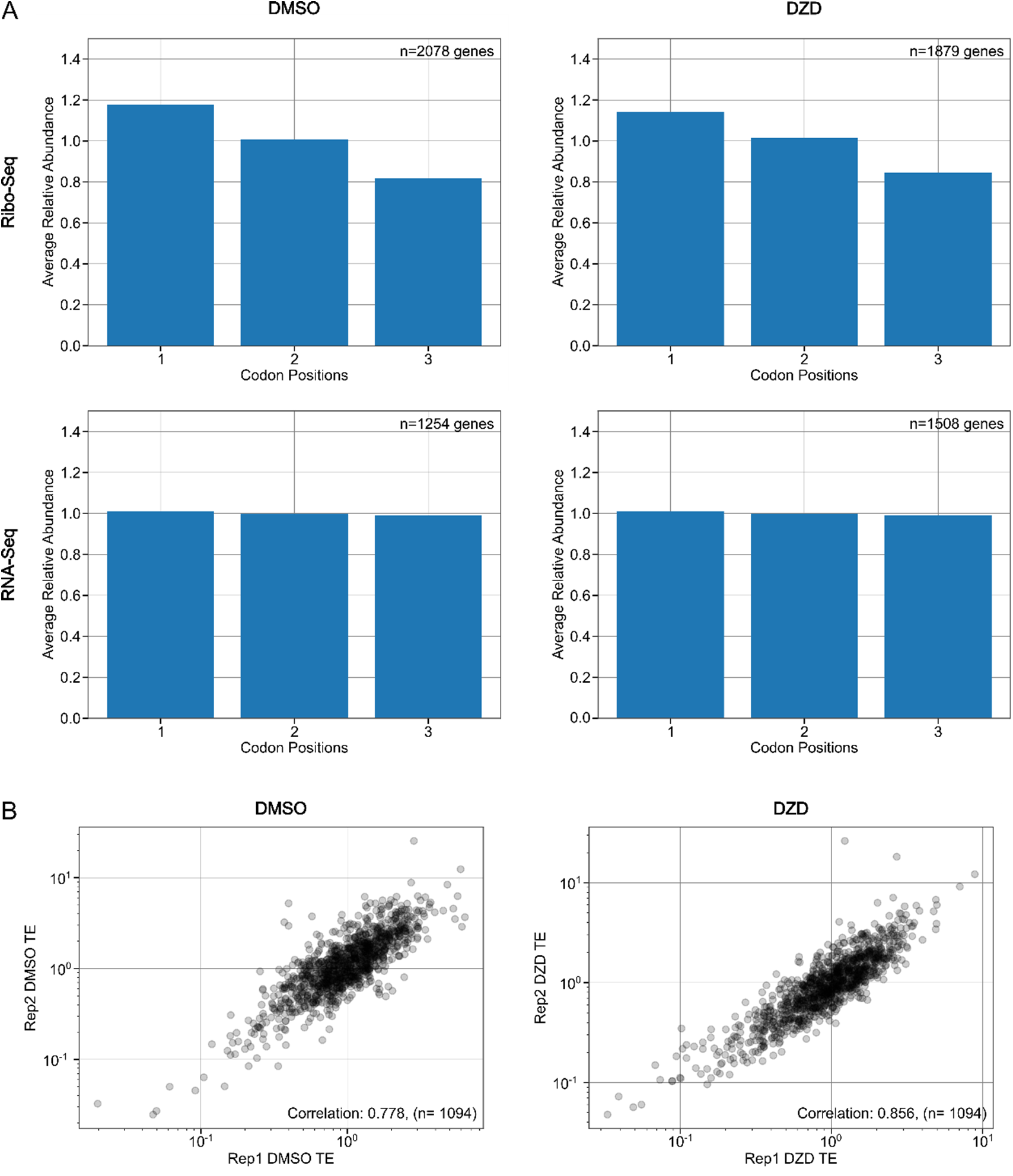
Ribosome Profiling Processing After DZD Treatment. **A)** Periodicity analysis of profiling data following 2.5 min treatment with DMSO or 10x MIC DZD. Top: Ribosome profiling. Bottom: RNA-seq. **B)** Genewise correlation between replicates. Changes in translation efficiency (RPKM Ribosome profiling/RPKM RNA-seq) for individual genes between experimental replicates following DMSO (left) or DZD (right) treatment.

**Extended Data Fig. 4.**
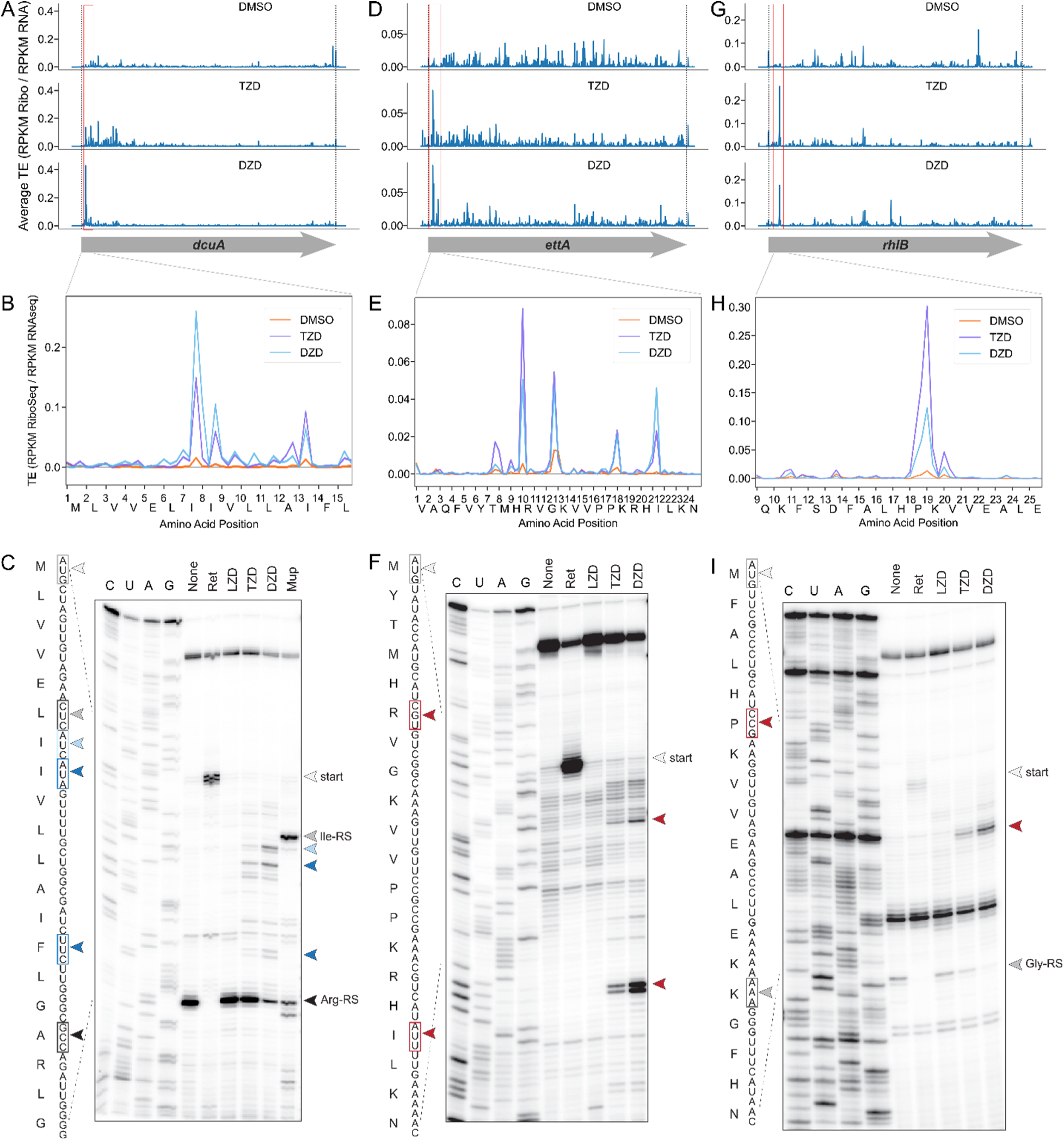
Additional validation of Ile(−1) and His(−1) stalling performed on genes *dcuA*. (**A-C**), *ettA* (D-F), and *rhlB* (G-I). **A, D, G)** Ribosome profiling traces across the relevant open reading frames after TZD or DZD treatment. Red boxes highlight prominent drug-induced peaks. **B, E, H)** Zoom into the most prominent stalling peaks for each gene. **C, F, I)** Toeprinting analysis in the presence of LZD, TZD, or DZD. Stalling by retapamulin (ret) at the start codon is shown in light gray, tRNA-synthetase inhibitors in dark gray or black, Ile(−1) in blue, Leu(−1) in light blue, and His(−1) in red.

**Extended Data Fig. 5.**
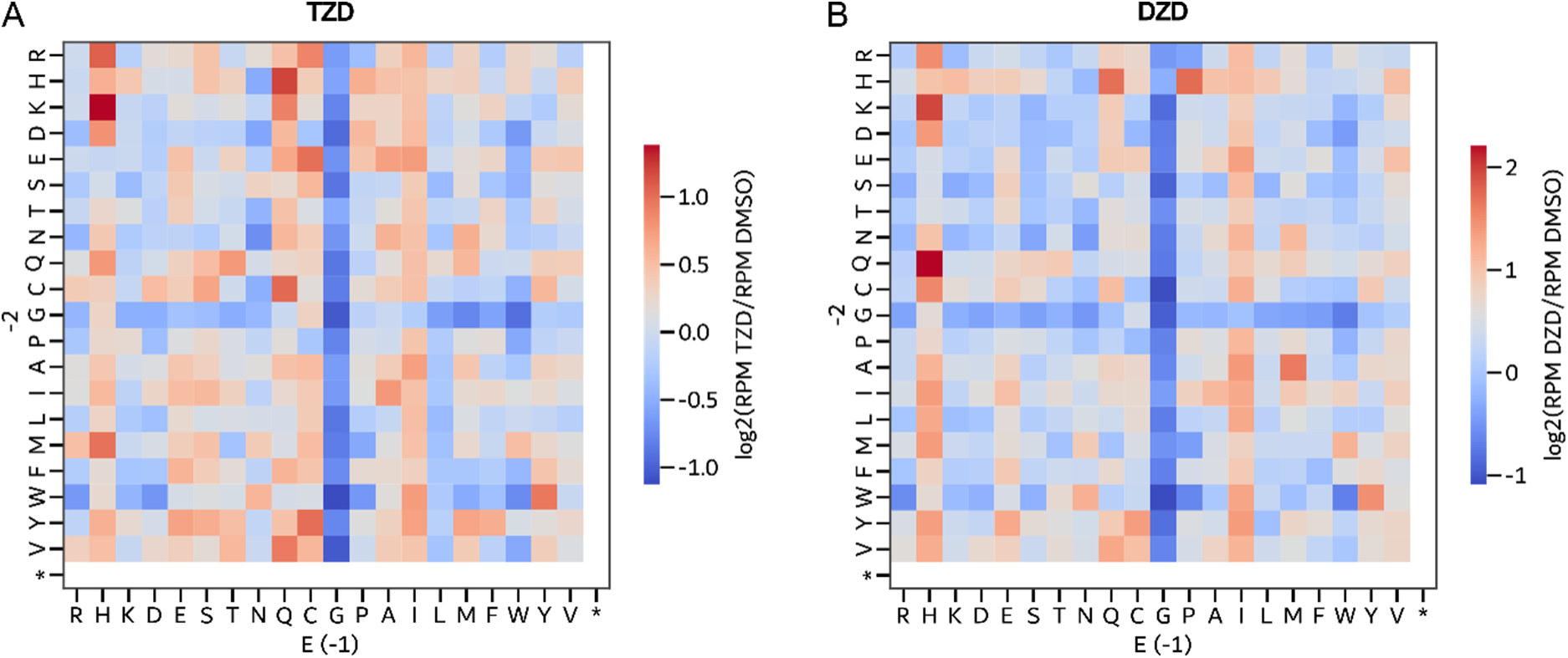
Analysis of extended stalling motifs in ribosome profiling data transcriptome-wide following treatment with A) TZD or B) DZD. Log2 fold change in reads per million compared to DMSO control was averaged across all instances of each –2:-1 amino acid pairing within the nascent chain in order to evaluate pairwise staling preferences.

**Extended Data Fig. 6.**
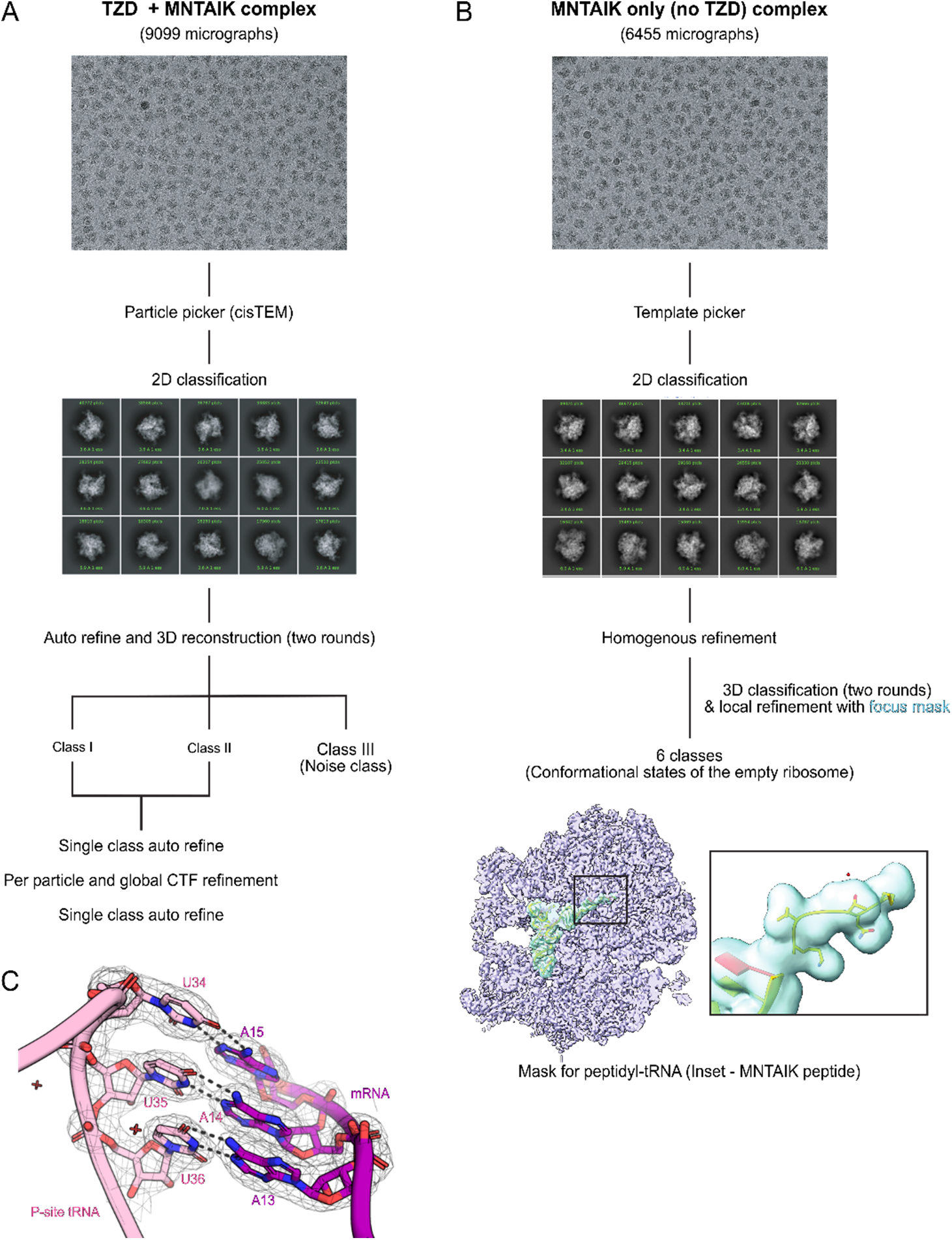
Processing workflow & classification tree for each map. **A)** For TZD-MNTAIK SRC: Data processing workflow in cisTEM for generating a consensus volume. After CTF estimation, particle picking, and 2D classification, 3D auto-refinement was performed for three classes, one of which was a junk class. The remaining two classes were identical. As a result, they were combined, followed by a round of single-class auto-refinement and CTF refinement. **B)** For MNTAIK-only complex: All the steps till homogeneous refinement were identical to the cisTEM workflow except the usage of the template picker in cryoSPARC as opposed to the automatic particle picker in cisTEM. 3D classification was performed after homogeneous refinement, using 6 classes with a class similarity score of 0.7. This was followed by focused refinement using a mask for P-site peptidyl-tRNA (Inset-Cyan) **C)** Density for the decoding center of TZD-MNTAIK SRC at 4.0σ. Nucleotides of P-site tRNA are shown in pink, and those of the mRNA are shown in purple. Dashed lines indicate hydrogen bonds.

**Extended Data Fig. 7.**
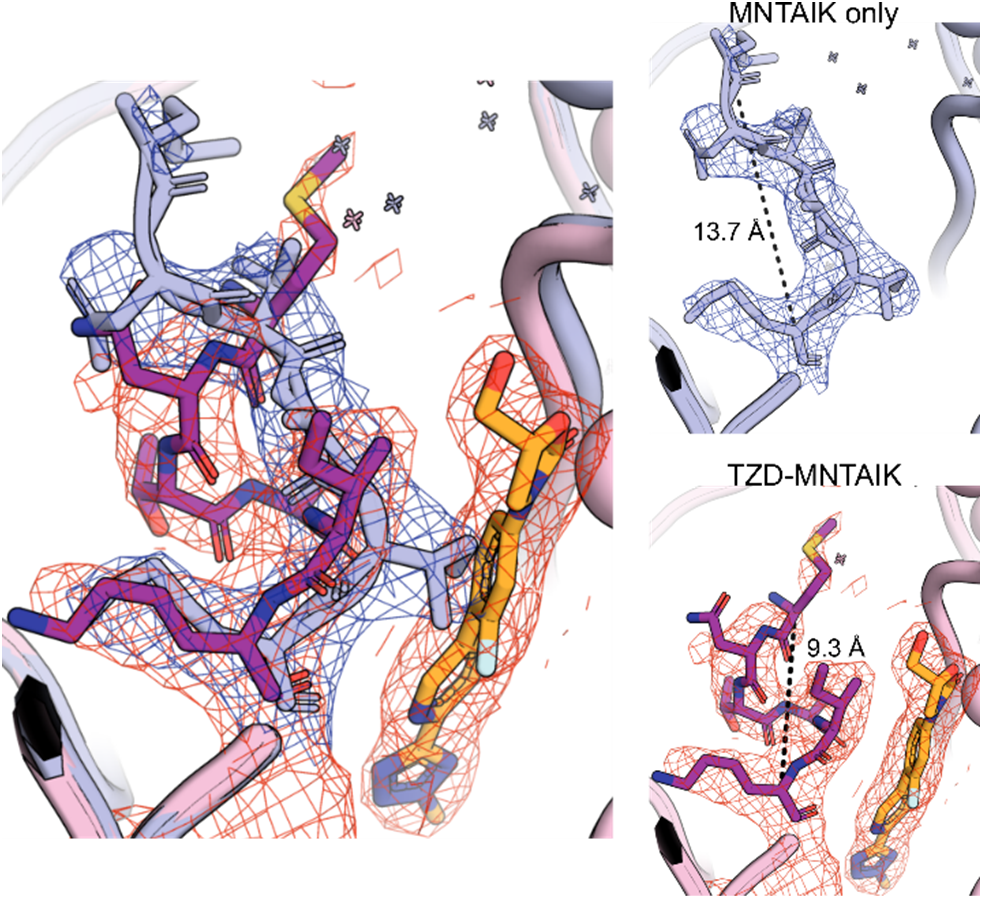
Path comparison of nascent peptides in drug-free and drug-bound ribosomes. Overlay of densities for MNTAIK in the TZD-SRC (peptide in magenta with red density) and the drug-free MNTAIK RNC (lavender, blue density). Inlay: measurement of peptide length from Met(−5) Cα to Lys(0) Cα, shown in black dashed lines, for each complex.

**Extended Data Fig. 8.**
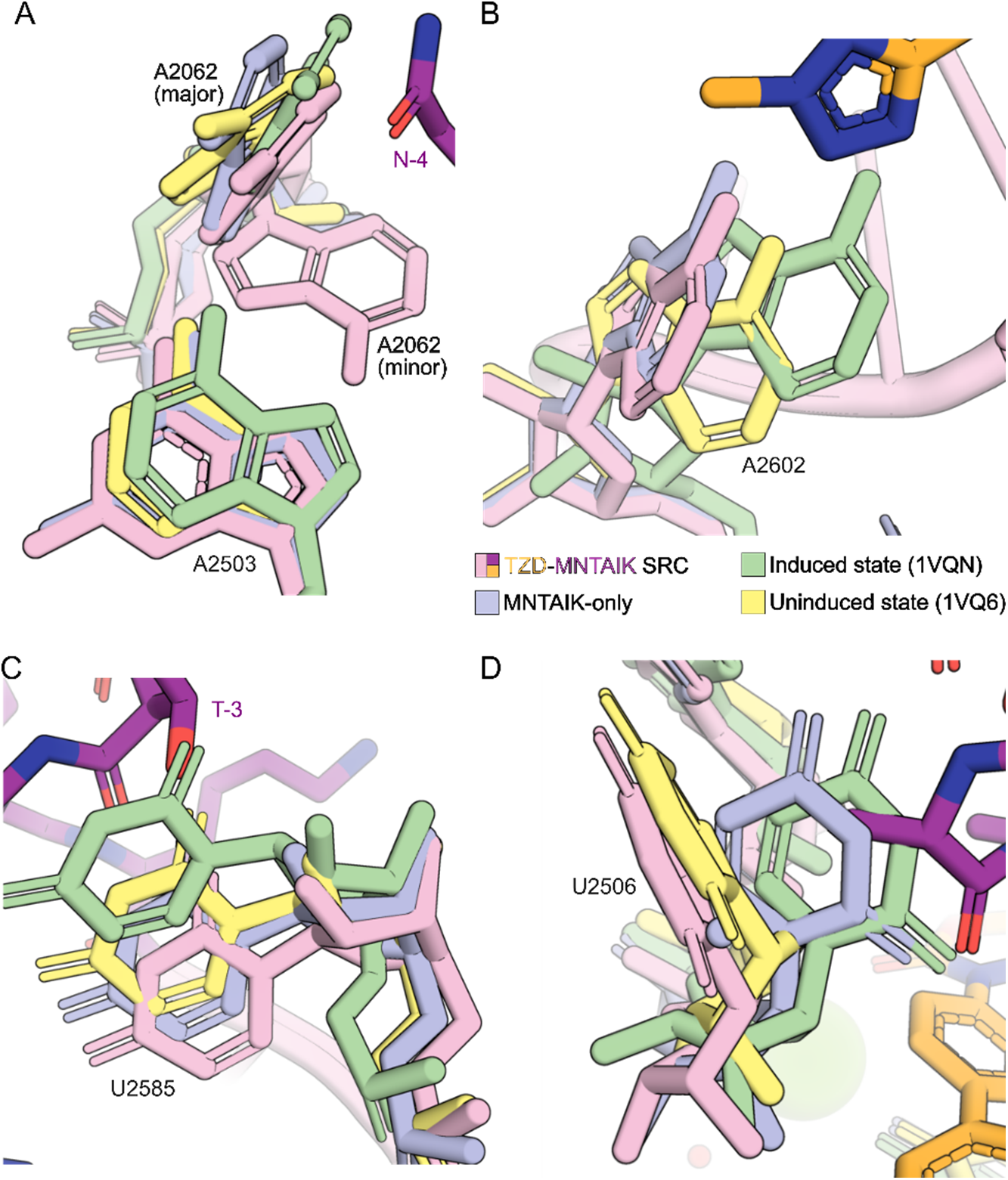
Nucleotide positions compared to a catalytically induced and uninduced state of the ribosome. TZD-MNTAIK SRC (rRNA: pink, peptide: magenta, TZD: gold) and MNTAIK-only RNC (lavender) compared to the catalytically favored (Induced state; green, PDB 1VQN) and catalytically nonproductive (Uninduced state; yellow, PDB 1VQ6) states of the *H. marismortui* ribosome^37^. **A)** The major conformation of A2062 in the TZD-SRC overlays with both the induced and uninduced conformations, while the minor conformation is distinct. **B)** Orientation of A2602 is distinct from those of the *H. mar* structures. **C)** Position of U2585 in TZD-MNTAIK SRC and MNTAIK-alone RNC is nearer to that of the uninduced conformation. **D)** Position of U2506 in the TZD SRC mimics that of the uninduced state, while U2506 conformation in the MNTAIK-only SRC is more similar to the induced state.

**Extended Data Fig. 9.**
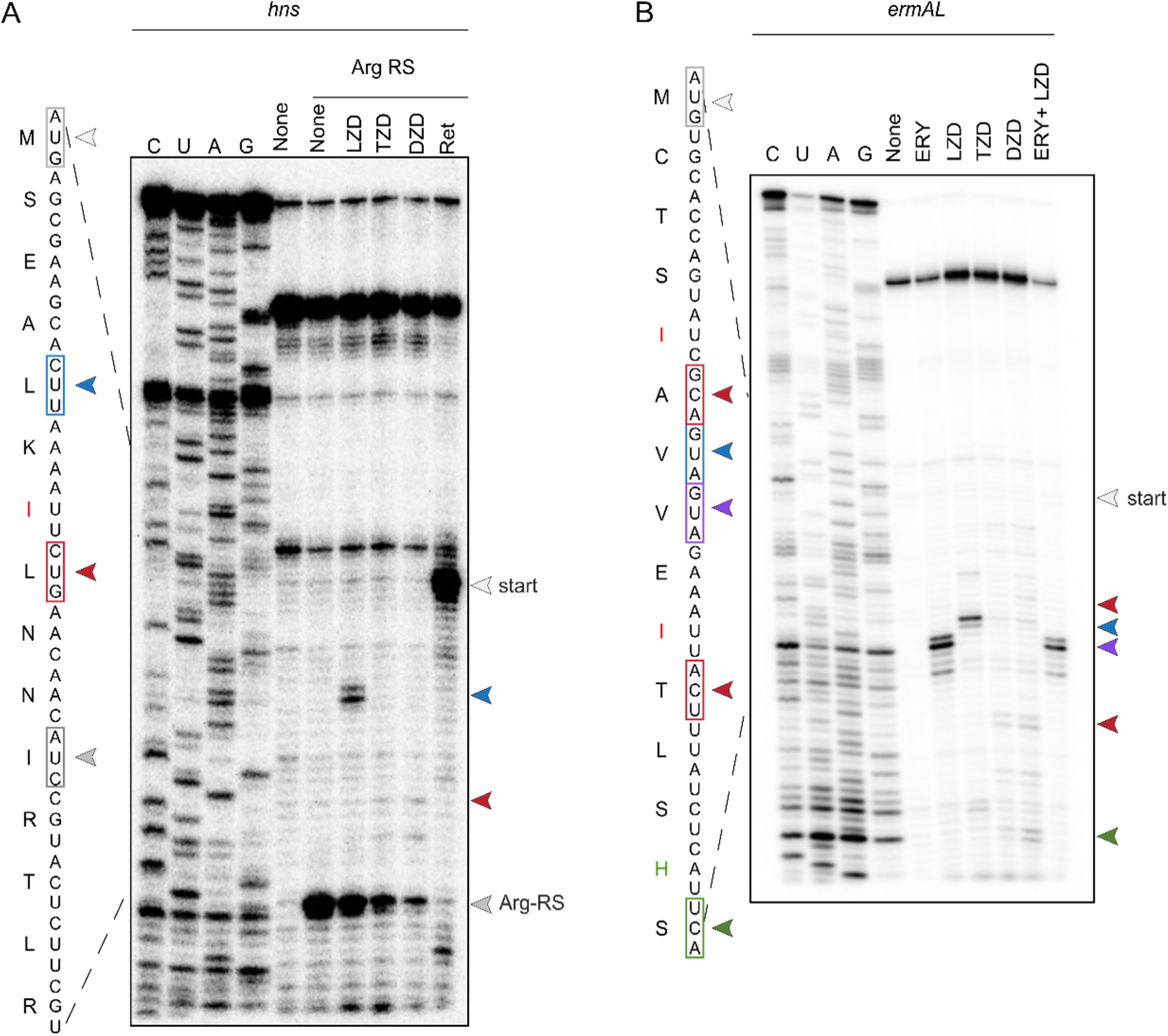
Stalling does not occur at all Ile(−1)–containing sites. Toeprinting on mRNA templates derived from *hns* (A) or Erm-resistance leader peptide *ermAL* (B) in the presence of LZD, TZD, or DZD. Stalling at start codon by retapamulin is shown in light gray, by Arg-RS at Arg(+1) in dark gray, and by LZD at Ala(−1) in blue. **A)** No stalling is observed for the oxazolidinones with Ile(−1) on the *hns* transcript (red). **B)** Little to no stalling is visible at any of the 3 putative TZD and DZD stall sites within *ErmAL* [red: Ile(−1); green: His(−1)]. Erythromycin-mediated stalling (purple) is observed at the canonical site for induction of downstream Erm-resistance genes^67^. All compounds were used at 50 μM.

## DATA AVAILABILITY

Atomic coordinates for all the structures presented here have been deposited in the Protein Data Bank and EMBD with the following accession numbers: TZD-MNTAIK-SRC: PDB ID 10ZD, EMDB-75563 and MNTAIK-only RNC: PDB ID 10ZE, EMDB-75564. The following datasets were used to generate the initial volumes for the presented structures: wild-type *E.coli* 70S ribosome subunit, PDB ID 7K00 (without the tRNAs).

## ACKNOWLEDGEMENTS

We thank D. Bulkley and G. Gilbert for technical support at the UCSF Center for Advanced CryoEM, which is supported by the National Institutes of Health (S10OD020054 and S10OD021741) and the Howard Hughes Medical Institute (HHMI). Sequencing was performed at the UCSF CAT, supported by UCSF PBBR, RRP IMIA, and NIH 1S10OD028511-01 grants. We acknowledge support from NIAID (R01AI137270 to D.G.F. and F31AI181568 to J.I.K.), NIGMS (R35GM145238 to J.S.F and R35GM127134 to A.S.M.), and W.M. Keck Foundation Medical Research Grant (to J.S.F. and D.G.F.).

## Notes

### Competing Interest Statement

DGF and JSF are co-founders of Interdict Bio.

