## Supplemental Information for "Structural modification of oxazolidinone antibiotics alters nascent peptide stalling preference and peptide trajectory through the ribosome"

AUTHORS & AFFILIATIONS

Jordan I. Kleinman^1,2^, Tushar Raskar^3,4^, Dorota Klepacki^5,6^, Teresa Szal^5,6^, Nora Vázquez-Laslop^5,6^, Alexander S. Mankin^5,6^, James S. Fraser^3,4,*^, Danica Galonić Fujimori^1,4,7,*^

^1^Department of Cellular and Molecular Pharmacology, University of California San Francisco, San Francisco, CA, USA.

^2^Chemistry and Chemical Biology Graduate Program, University of California San Francisco, San Francisco, CA, USA

^3^Department of Bioengineering and Therapeutic Sciences, University of California San Francisco, San Francisco, CA, USA.

^4^Quantitative Biosciences Institute, University of California, San Francisco, San Francisco, CA 94143, USA.

^5^Department of Pharmaceutical Sciences, University of Illinois at Chicago, Chicago, IL, USA. ^6^Center for Biomolecular Sciences, University of Illinois at Chicago, Chicago, IL, USA.

^7^Department of Pharmaceutical Chemistry, University of California San Francisco, San Francisco, CA, USA.

**Supplementary Table 1. Primer Sequences used in this study.**

| **Supplementary Table X: Primer sequences** | | |
| --- | --- | --- |
| **Name** | **Sequence** | **Purpose** |
| NV1 | GGTTATAATGAATTTTGCTTATTAAC | Toeprinting |
| FKAFK | GCCCCGGTAAGGAAATAAAAATGTTCAAAGCATTCAAAAACATCATACGTACTCGTACTC | Toeprinting |
| FKAFK-Rev | GGTTATAATGAATTTTGCTTATTAACCTTGCCTGCGCTTAAAGAGTACGAGTACGTATGATGT | Toeprinting |
| FWD | ATTAATACGACTCACTATAGGGCAACCTAAAACTTACACACGCCCCGGTAAGGAAATAAAAAT | Toeprinting |
| T7-IR-fwd | TAATACGACTCACTATAGGGCTTAAGTATAAGGAGGAAAACAT | Toeprinting |
| IR-rhlB | GTATAAGGAGGAAAACATATGTTCGCCCTGCATCCGAAGG | Toeprinting |
| NV1-rhlB-rev | GGTTATAATGAATTTTGCTTATTAACCGCCAGCGTCAGCGGAAGGGC | Toeprinting |
| IR-ettA | GTATAAGGAGGAAAACATATGTATACCATGCATCGTGTCGGC | Toeprinting |
| NV1-ettA-rev | GGTTATAATGAATTTTGCTTATTAACGACACCAATTTTTGCCCCAG | Toeprinting |
| IR-ispH | GTATAAGGAGGAAAACATATGTATGTCCGTCACGAAGTGGTAC | Toeprinting |
| NV1-ispH-rev | GGTTATAATGAATTTTGCTTATTAACGTCCGGTACTTCGCTAATC | Toeprinting |
| IR-typA | GTATAAGGAGGAAAACATATGAACACCGCTATCAAATGGAA | Toeprinting |
| NV1-typA-rev | GGTTATAATGAATTTTGCTTATTAACCATGGACATTACACGTTCAAC | Toeprinting |
| IR-dcuA | GTATAAGGAGGAAAACATATGCTAGTTGTAGAACTCATC | Toeprinting |
| NV1-dcuA-rev | GGTTATAATGAATTTTGCTTATTAACCAGCACCCCCAATCCGCCTG | Toeprinting |
| SC-15 | rArUrGrUrArCrArCrGrGrArGrUrCrG | RiboSeq size selection |
| SC-45 | rArUrGrUrArCrArCrGrGrArGrUrCrGrArCrCrCrGrCrArArCrGrCrGrArUrGrUrArCrArCrGrGrArGrUrCrGrArCrC | RiboSeq size selection |
| Universal miRNA cloning linker (NEB S1315S) | 5′ rAppCTGTAGGCACCATCAAT–NH2 3′ | Ligation linker |
| Ni-Ni-9 | /5Phos/AGATCGGAAGAGCGTCGTGTAGGGAAAGAGTGTAGATCTCGGTGGTCGC/iSp18/CACTCA/iSp18/TTCAGACGTGTGCTCTTCCGATCTATTGATGGTGCCTACAG | Reverse transcription |
| b-rRNA-1 | /5Biosg/TCATCTCCGGGGGTAGAGCACTGTTTCG | rRNA depletion |
| b-rRNA-2 | /5Biosg/GGCTAAACCATGCACCGAAGCTGCGGCAG | rRNA depletion |
| b-rRNA-3 | /5Biosg/AAGGCTGAGGCGTGATGACGAGGCACT | rRNA depletion |
| b-rRNA-4 | /5Biosg/CGGTGCTGAAGCAACAAATGCCCTGCTT | rRNA depletion |
| b-rRNA-5 | /5Biosg/CTGCTTCCAGGAAAAGCCTCTAAGCATCAGG | rRNA depletion |
| P5 | AATGATACGGCGACCACCGAGATCTACAC | NGS amplification |
| P7_i23 | CAAGCAGAAGACGGCATACGAGATCCACTCGTGACTGGAGTTCAGACGTGTGCTCTTCCG | NGS amplification |
| P7_i24 | CAAGCAGAAGACGGCATACGAGATGCTACCGTGACTGGAGTTCAGACGTGTGCTCTTCCG | NGS amplification |
| P7_i26 | CAAGCAGAAGACGGCATACGAGATGCTCATGTGACTGGAGTTCAGACGTGTGCTCTTCCG | NGS amplification |
| P7_i27 | CAAGCAGAAGACGGCATACGAGATAGGAATGTGACTGGAGTTCAGACGTGTGCTCTTCCG | NGS amplification |
| P7_i28 | CAAGCAGAAGACGGCATACGAGATCTTTTGGTGACTGGAGTTCAGACGTGTGCTCTTCCG | NGS amplification |
| P7_i31 | CAAGCAGAAGACGGCATACGAGATATCGTGGTGACTGGAGTTCAGACGTGTGCTCTTCCG | NGS amplification |
| P7_i32 | CAAGCAGAAGACGGCATACGAGATTGAGTGGTGACTGGAGTTCAGACGTGTGCTCTTCCG | NGS amplification |
| MNTAIK_R | TTTGATAGCGGTGTTCAT | SRC-rev-short |
| MNTAIK.W_R | CCATTTGATAGCGGTGTTC | SRC-rev-short |
| SD_MNTAIK_R | TTTGATAGCGGTGTTCATTTTTATTTCCTTACCGGGGCGTGTGTAAGTTTTAG | SRC-rev-long |
| SD_MNTAIK.W_R | CCATTTGATAGCGGTGTTCATTTTTATTTCCTTACCGGGGCGTGTGTAAGTTTTAG | SRC-rev-long |
| T7 | ATTAATACGACTCACTATAGGG | Toeprinting, SRC-fwd-short |
| T7_fwd_long | ATTAATACGACTCACTATAGGGCAACCTAAAACTTACACACGCCCCG | SRC-fwd-long |
